# RhoA within myofibers controls satellite cell microenvironment to allow hypertrophic growth

**DOI:** 10.1101/2021.01.18.426685

**Authors:** Chiara Noviello, Kassandra Kobon, Léa Delivry, Thomas Guilbert, Francis Julienne, Pascal Maire, Voahangy Randrianarison-Huetz, Athanassia Sotiropoulos

## Abstract

Adult skeletal muscle is a plastic tissue that can adapt its size to workload. Here, we show that RhoA within myofibers is needed for overload-induced hypertrophy by controlling satellite cell fusion to the growing myofibers without affecting protein synthesis. At the molecular level, we demonstrate that, in response to increased workload, RhoA controls in a cell autonomous manner Erk1/2 activation and the expressions of extracellular matrix (ECM) regulators such as *Mmp9/Mmp13/Adam8* and of macrophage chemo-attractants such as *Ccl3*/*Cx3cl1.* Their decreased expression in RhoA mutant is associated with ECM and fibrillar collagen disorganization and lower macrophage infiltration. Moreover, Mmps inhibition and macrophage depletion in controls phenocopied the lack of growth of RhoA mutants. These findings unravel the implication of RhoA within myofibers, in response to increase load, in the building of a permissive microenvironment for muscle growth and for satellite cell accretion through ECM remodeling and inflammatory cell recruitment.

## Introduction

Skeletal muscle is a highly plastic tissue among the most abundant in the vertebrate body. The cellular muscle unit is the myofiber that is mainly composed of sarcomeric proteins with contractile properties. Physiological demands such as exercise or functional overload (OV) lead to an increase of muscle mass due to a hypertrophic growth of myofibers. In the adult the two major mechanisms, related to muscle growth and to sarcoplasmic volume enlargement are: i) protein synthesis increase to add new contractile filaments to pre-existing sarcomere units, and ii) the fusion of new nuclei provided by resident muscle stem cells, the satellite cells (SCs). In response to increased workload, SCs exit the quiescent state, proliferate, differentiate and subsequently fuse to growing myofibers (Fukada et al., 2020).

Furthermore, at the cellular level, skeletal muscle tissue homeostasis requires the coordinated function of other cell types present in the muscle itself. During the regenerative process, a network of interactions of SCs with different cell types including macrophages, endothelial cells and fibro/adipogenic progenitors (FAPs) orchestrates muscle regeneration. Indeed, macrophages play various sequential roles during skeletal muscle regeneration. At the early stages of injury, pro-inflammatory macrophages stimulate SC proliferation and later on, after their conversion into anti-inflammatory macrophages, they support SC differentiation and fusion into myofibers (Chazaud, 2020). However, the contribution of muscle resident cells other than SCs and myofibers to muscle hypertrophy and the role of muscle microenvironment remain poorly studied so far.

In addition, modifications of muscle tissue microenvironment, such as ECM remodeling, occurring upon increased workload have not been fully explored. The major roles of ECM are: i) to create tissue scaffold for vessels and nerves; ii) to contribute to the force transmission through the passive elastic response of muscle iii) to regulate dynamically cell functions through ECM-embedded molecules such as growth factors. In addition ECM provides a structural support to the integrity of SC niche, physically separating the stem cell pool from other tissue resident cells, and can thus play an important role in SC accretion during hypertrophy. Among the proteins mostly present in ECM are Collagens, Fibronectin (Fn1) and TenascinC (TnC) that are crucial components of SC niche regulating their self-renewal (Bentzinger et al., 2013; Urciuolo et al., 2013) and their regenerative potential (Tierney et al., 2016).

The major enzymes responsible for the physiological breakdown of ECM are matrix metalloproteinases (Mmps). One of the best characterized Mmp in SCs and muscle tissue is Mmp9 (Chenette et al., 2016; Rayagiri et al., 2018). *Mmp9* deletion is deleterious to muscle mass (Mehan et al., 2011) while its constitutive muscle-specific over-expression promotes growth (Dahiya et al., 2011) supporting the notion that a precise regulation of muscle ECM is necessary during periods of adaptation to ensure optimal muscle growth. Nevertheless, how ECM is remodeled and regulated upon increased workload remains unclear.

Ras homolog family member A (RhoA) is a small GTPase protein that oscillates between GTP bound and GDP bound states, regulating a wide spectrum of cellular functions. RhoA controls contractility, actin polymerization and actin cytoskeleton organization. RhoA is an important player of mechanotransduction that translates physical forces into biochemical signaling pathways and transmits the signal in the nucleus, activating specific transcription factors (Lessey et al., 2012; Burridge et al., 2019). Among them, Serum response factor (Srf) activity is regulated by RhoA through the control of actin dynamics (Sotiropoulos at al, 1999). In muscle tissue, mRNA and protein expressions of RhoA have been shown to increase upon chronic functional OV and hypertrophy-stimulating resistance training in rat and human, respectively (McClung et al., 2003). *In vitro*, cyclic stretch activated RhoA in cultured myotubes (Zhang et al., 2007). However, the functional role of RhoA in skeletal muscle physiology and in muscle mass regulation has not been investigated.

In this study, we assessed the role played by RhoA within myofibers during skeletal muscle hypertrophy by inducing compensatory hypertrophy of *plantaris* muscles harbouring a conditional and inducible deletion of *RhoA* in myofibers. We showed compromised hypertrophic growth in absence of RhoA. The impaired growth was not associated with a protein synthesis defect but rather with altered SC fusion to the growing myofibers. We showed that the most downregulated genes in OV mutant (Mut) muscles compared to controls (Ctl) are those involved in ECM remodeling, like *Mmp9/13* and *Adam8*. Furthermore, a decreased expression of these genes in Mut is associated with un-degraded ECM proteins and fibrillar collagen disorganization. Inhibition of Mmps activity affected muscle growth by preventing SC recruitment and thus phenocopying RhoA mutants. Finally, we showed that the expression of potent chemo-attracting chemokines (*Cx3Cl1* and *Ccl3*) is reduced in *RhoA*-deleted myofibers upon OV and is associated with diminution of macrophage infiltration which functional necessity was demonstrated by depletion experiments.

Altogether our data highlighted a new role of RhoA within myofibers in the regulation of muscle microenvironment upon OV. By modulating ECM remodeling and inflammation, RhoA may allow the constitution of a correct SC niche that will affect SC behaviour and thus permits a correct hypertrophic growth.

## Results

### RhoA is required in myofibers for overload-induced hypertrophy

In order to investigate the signaling pathways functionally involved in the control of adult skeletal muscle growth in response to increased load, we performed a transcriptomic study of genes expressed in *plantaris* muscles at the basal state (Sham Operated, SO) and 1 week after overload-induced hypertrophy (OV 1wk). We identified genes differentially expressed before and after OV (fold change>1.2 and pvalue<0.05). Analysis of the Canonical Pathways using Ingenuity Pathway Analysis (IPA) pointed out RhoA signaling among the first 10 pathways ranked by their significance with predicted activation (Z-score>2) (Table S1). We thus hypothesized that RhoA signaling could have a crucial role during skeletal muscle hypertrophy.

To examine the contribution of RhoA during muscle hypertrophic growth, we generated a mouse model in which *RhoA* is deleted in a specific and inducible manner only in the adult myofiber compartment by injecting tamoxifen (Tmx) to *HSA-Cre^ERT2^:Rho^flox/flox^* mice (refered as mutant) and performed OV-induced hypertrophy of the *plantaris* muscle (Figure 1A). At the steady state (SO), a significant 60% RhoA loss was achieved at the protein and transcript levels in different muscles including *gastrocnemius, tibialis anterior* and *plantaris* (Figure 1B, 1C). Because different cell types are present in the muscles and myofiber myonuclei represent roughly 50% of the total nuclei (Dos Santos et al., 2020), we checked the expression of *RhoA* transcripts in isolated control (Ctl) and mutant (Mut) *plantaris* single myofibers. We showed an efficient 85% mRNA decrease upon Tmx treatment (Figure 1D, SO condition). No difference of muscle weight and myofiber Cross Section Area (CSA) was observed before hypertrophy between Ctl and Mut *plantaris* (Figure 1E, 1F, 1G). Following OV, *RhoA* expression increased at the onset of the hypertrophy process (1wk after surgery) in muscle tissue and isolated myofibers (Figure 1D) in agreement with the increased *RhoA* expression observed in overloaded rat muscles (McClung et al., 2003). Three weeks post OV, *plantaris* muscle mass and CSA were increased in Ctl as compared to unloaded muscles (SO Ctl). However, the extent of hypertrophic growth was decreased in Mut *plantaris* muscles. They displayed a 15% increase in muscle weight (after 3wk OV) instead of 30% in Ctl and no significant increase of their CSA (Figure 1E, 1F, 1G). In addition, the increase in muscle mass was mainly caused by myofiber hypertrophy as the total number of myofibers did not vary after OV (Figure 1H). These data show that RhoA in the myofibers is necessary for optimal myofiber OV-induced hypertrophy and that its absence causes muscle growth defect.

**Figure 1.**
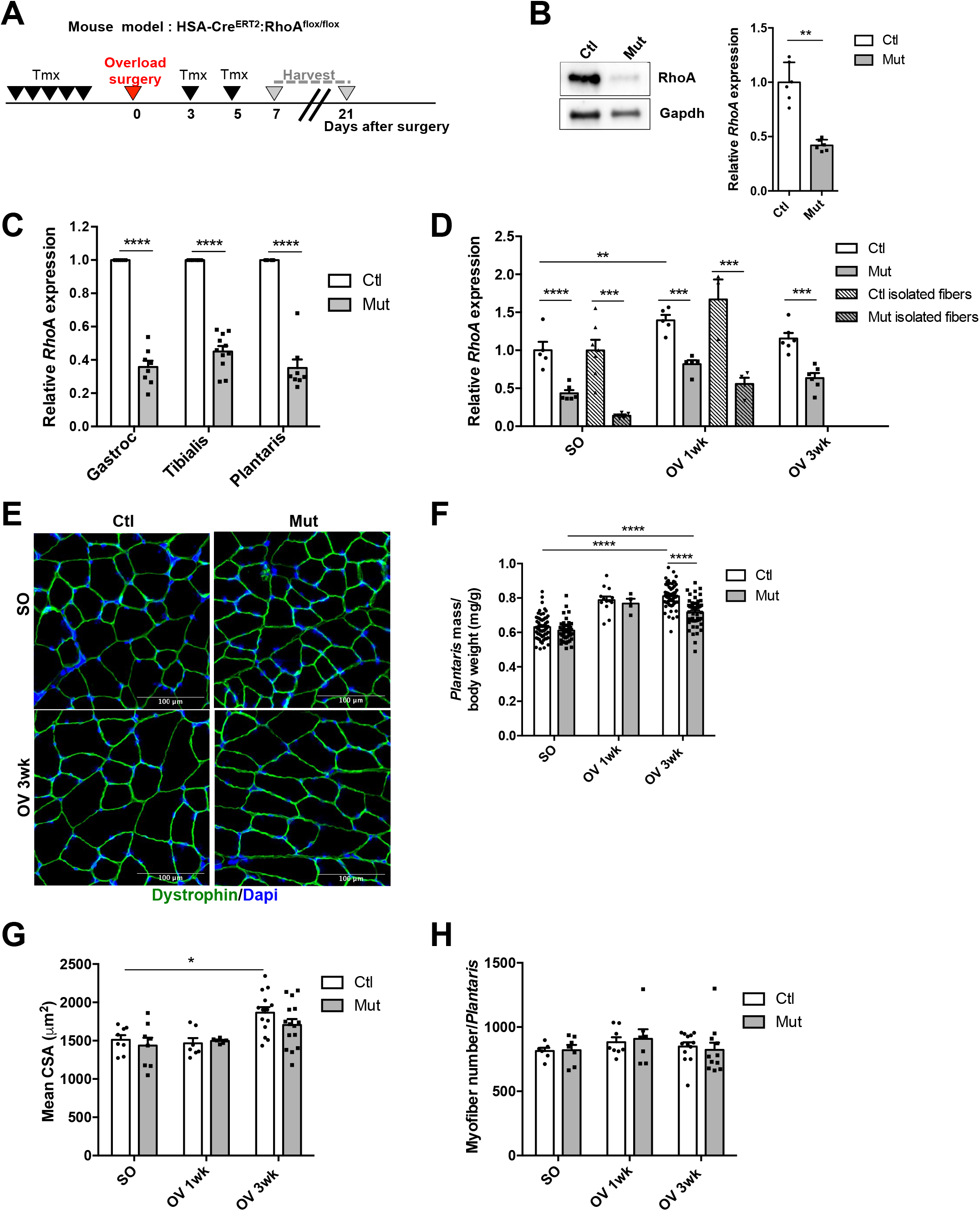
RhoA loss within myofibers impairs overload-induced hypertrophy. **(A)** RhoA mutant (Mut) mice were injected with Tmx 1 week (wk) before OV procedure. *Plantaris* muscles were isolated 1 and 3wk after surgery. **(B)** RhoA protein was analysed by Western blot in Ctl and Mut *plantaris* muscles. Gapdh was used as a loading control. **(C)** Analysis of *RhoA* mRNA expression by RT-qPCR in Ctl and Mut *gastrocnemius, tibialis anterior* and *plantaris* muscles. Data were normalized by *Hmbs* expression and relative to Ctl (n=6-11). **(D)** Analysis of *RhoA* mRNA expression by RT-qPCR in Ctl and Mut *plantaris* muscles or isolated myofibers before (SO) and after 1 and 3wk OV. Data were normalized by *Hmbs* expression and relative to Ctl SO (n=3-7). **(E)** *Plantaris* sections immunostained for Dystrophin (green) and nuclear staining with DAPI for Ctl and Mut before (SO) and after 3wk OV. **(F)** Ratio of *plantaris* mass (mg) to body weight (g) before (SO) and after 1 and 3wk OV in Ctl and Mut (n=5-20). **(G)** Mean CSA (μm^2^) before (SO) and after 1 and 3wk OV in Ctl and Mut (n=6-14). **(H)** Mean myofiber number before (SO) and after 1 and 3wk OV in Ctl and Mut. Data and mean±SEM. *pvalue<0.05, ***pvalue<0.001, ****pvalue<0.0001.

### RhoA is dispensable for protein synthesis control upon hypertrophy

Hypertrophic growth is associated with an increase of protein synthesis and a decrease of protein degradation at the onset of compensatory hypertrophy procedure. Akt signaling plays a central role in this control by activating mTOR pathway and reducing FoxO-mediated atrogenes transcription (Sandri et al., 2004). So, we first tested whether the impaired OV-induced hypertrophy of Mut *plantaris* muscles could be attributed to impaired global protein synthesis. The rate of total protein synthesis was measured *in vivo* using the SUnSET technique that relies on the incorporation of Puromycin in newly translated proteins. We showed a similar augmentation in both Ctl and Mut muscles 1wk following OV with no significant difference between the two groups (Figure S1A). We then analysed Akt and p70S6K signaling and we observed no difference in Akt and p70S6K phosphorylations between Ctl and Mut, with a 6 fold and 2.5 fold increase upon OV respectively (Figure S1B). By phosphorylating FoxO transcription factors, Akt activation prevents their nuclear localisation and the transcription of their target genes, among which the atrogenes encoding MuRF1 and MAFbx ubiquitin ligases. In line with the activation of Akt, *MuRF1* and *MAFbx* expressions decreased in a similar manner in Ctl and Mut *plantaris* 1wk post OV and came back to basal levels after 3wk (Figure S1C). Altogether these data suggest that the absence of RhoA in the myofibers does not affect protein synthesis and degradation pathways upon OV-induced hypertrophy.

### Loss of RhoA in myofibers impairs SC behaviours and in particular their fusion

In addition to the increase of protein content, muscle compensatory hypertrophy relies on the accretion of new nuclei through the mobilization of SCs. We hypothesized that RhoA in the myofibers may control the behaviour of SCs under OV-induced hypertrophic conditions. We investigated whether RhoA loss in myofibers altered the number of SCs, expressing Pax7 and RhoA. Before OV, Pax7^+^ SC number was identical between Ctl and Mut (Figure 2A, SO). One week after OV, there was a significant increase in Pax7^+^ SCs in both genotypes, but the number of SCs was slightly lower in Mut muscles. We further assessed the proliferative potential of SCs by determining the number of SCs that entered S-Phase (*in vivo* EdU injection 24hr and 4hr prior the end of the experiment – Pax7^+^EdU^+^ cells). However, the number of Pax7^+^EdU^+^ proliferating SCs and the proportion of proliferating SCs (Pax7^+^EdU^+^/Pax7^+^ cells) were not significantly affected (Figure 2B and 2C). Overall, following an increased load, RhoA in myofibers may slightly affect the proliferative and/or the activation potential of SCs.

**Figure 2.**
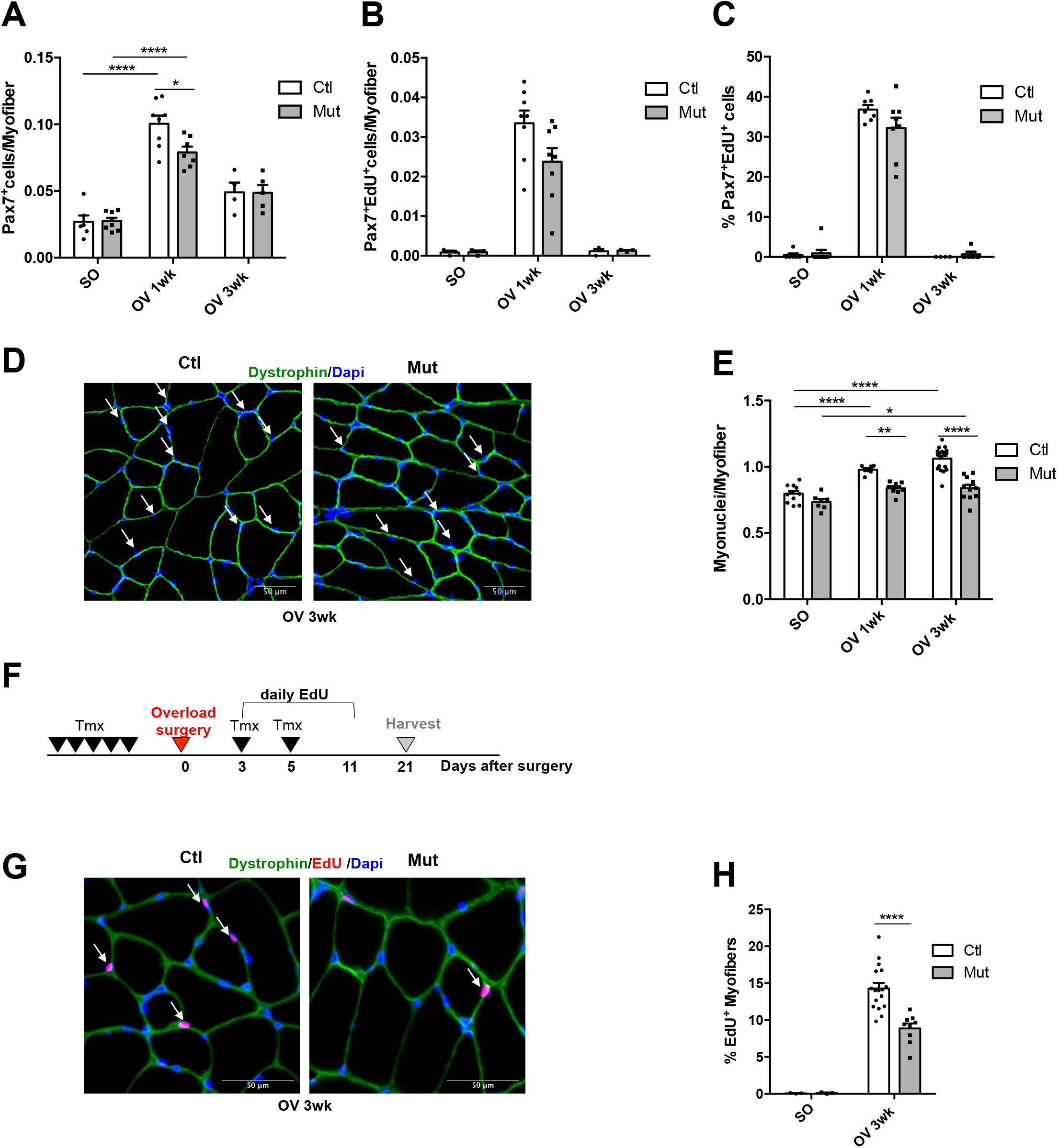
RhoA loss within myofibers impairs SC fusion. **(A)** Number of Pax7^+^ cells per myofiber (n=4-8) (A). **(B)** Number of Pax7^+^EdU^+^ cells per myofiber (n=3-8). **(C)** Percentage of Pax7^+^EdU^+^ among Pax7^+^ cells (n=4-8) in Ctl and Mut *plantaris* sections before (SO) and after 1 and 3wk OV. **(D)** *Plantaris* sections immunostained for Dystrophin (green) and nuclear staining with DAPI for Ctl and Mut after 3wk of OV. White arrows indicate myonuclei within Dystrophin^+^ sarcolemma. Number of myonuclei within sarcolemma per myofiber before (SO) and after 1 and 3wk OV (n=7-19). **(F)** Ctl and Mut were injected daily with EdU from day 3 to 11 post OV. *Plantaris* were isolated after 3wk OV. **(G)** *Plantaris* sections immunostained for Dystrophin (green), EdU (red) and nuclear staining with DAPI for Ctl and Mut mice before (SO) and after 3wk OV. **(H)** Percentage of EdU^+^ myofibers in Ctl and Mut *plantaris* sections before (SO) and after 3wk OV (n=9-14). Data and mean±SEM. *pvalue<0.05, **pvalue<0.01, ****pvalue<0.0001.

We next investigated whether *RhoA* deletion in the myofiber could affect SCs myogenic differentiation potential and their fusion to the Mut myofibers. Upon OV, *Myogenin* transcript levels did not differ between Ctl and Mut *plantaris*, with a sharp increase 1wk post OV that came back to SO levels after 3wk (Figure S2). Thus the loss of RhoA within myofibers did not affect the engagement of SCs in myogenic differentiation. Then, the fusion potential of SCs to the growing myofibers was assessed by counting the number of myonuclei within the Dystrophin-labeled myofiber sarcolemma (Figure 2D). The number of myonuclei per myofiber was similar before OV between Ctl and Mut and significantly increased 1 and 3wk after OV in Ctl and to a lesser extend in Mut muscles 3wk after OV (Figure 2E). Importantly, in Mut *plantaris*, the number of myonuclei was significantly decreased as compared to Ctl 1 and 3wk after OV (Figure 2E).

To further assert the impaired fusion capacities of SCs to the Mut myofibers *in vivo*, we chronically injected mice subjected to compensatory hypertrophy with EdU and we tracked EdU^+^ nuclei incorporated into growing myofibers (Figure 2F, 2G). Three weeks after OV, the percentage of EdU^+^ myofibers was significantly reduced in Mut compared to Ctl (Figure 2H). These data suggest that within myofibers RhoA is needed for an efficient mobilisation of SCs during hypertrophy by controlling their number and their recruitment to the growing myofibers. The altered behaviour of SCs in absence of RhoA could account, at least in part, for the impaired growth observed in mutant.

### RhoA control of SC behaviours is environment dependent

To investigate whether the altered behaviours of SCs, especially cell fusion, in *RhoA*-deleted muscles subjected to OV, are cell autonomous and unrelated to the local environment and/or mechanical stimuli, we set up an *in vitro* assay. This assay was designed to assess whether the absence of RhoA within myotubes (Adeno-Cre-mCherry transduced *RhoA^flox/flox^* myotubes) affects their fusion with RhoA-expressing mononucleated myocytes (expressing GFP) (Figure S3A, S3C). Surprisingly the lack of *RhoA* expression in myotubes (Figure S3B) did not alter the ability of myocytes to fuse to myotubes *in vitro* (dual-labeling) (Figure S3D). These data suggest that the defect on SC recruitment to *RhoA* mutant myofibers observed *in vivo* relies on muscle environment linked to the *in vivo* OV context, conditions that cannot be reproduced *in vitro*. This highlights the importance of mechanical cues and of *in vivo* microenvironment in the control of muscle compensatory growth. To further analyse how RhoA in the myofibers may control hypertrophic growth, we focused on *in vivo* studies.

### Mmps and chemokines expression is affected in absence of RhoA within myofibers

In order to decipher the molecular mechanisms underlying the growth defects of *RhoA*-deleted myofibers, we performed a genome wide microarray analysis of gene expression from Ctl and Mut *plantaris* muscles, before and 1wk after OV procedure. Gene expression profiling evidenced that a massive change occurred in muscles upon increased load, with more than 6000 genes up- and down-regulated in Ctl and in Mut muscles (SO vs OV 1wk; fold change>1.2; pvalue<0.05). Upon OV, 569 genes were differentially expressed between Ctl and Mut *plantaris* (Figure 3A). Among the 233 down-regulated genes in OV Mut muscles as compared to Ctl, we focused our attention on genes up-regulated in Ctl after hypertrophy (and to a lesser extend in mutants) (Figure 3B, highlighted by a red box). The 30 most differentially expressed genes of this category (Mut OV vs Ctl OV) are shown in Figure 4C. Among them, we identified genes encoding proteins involved in ECM remodeling, like *Mmp9* gelatinase, *Mmp13* collagenase and *Adam8* metalloproteinase, and important chemokines like *Cx3cl1* and *Ccl3* (Figure 3C). We confirmed by RT-qPCR that these genes were strongly and transiently expressed upon OV in Ctl while their expression was significantly reduced in Mut (Figure 3D).

**Figure 3.**
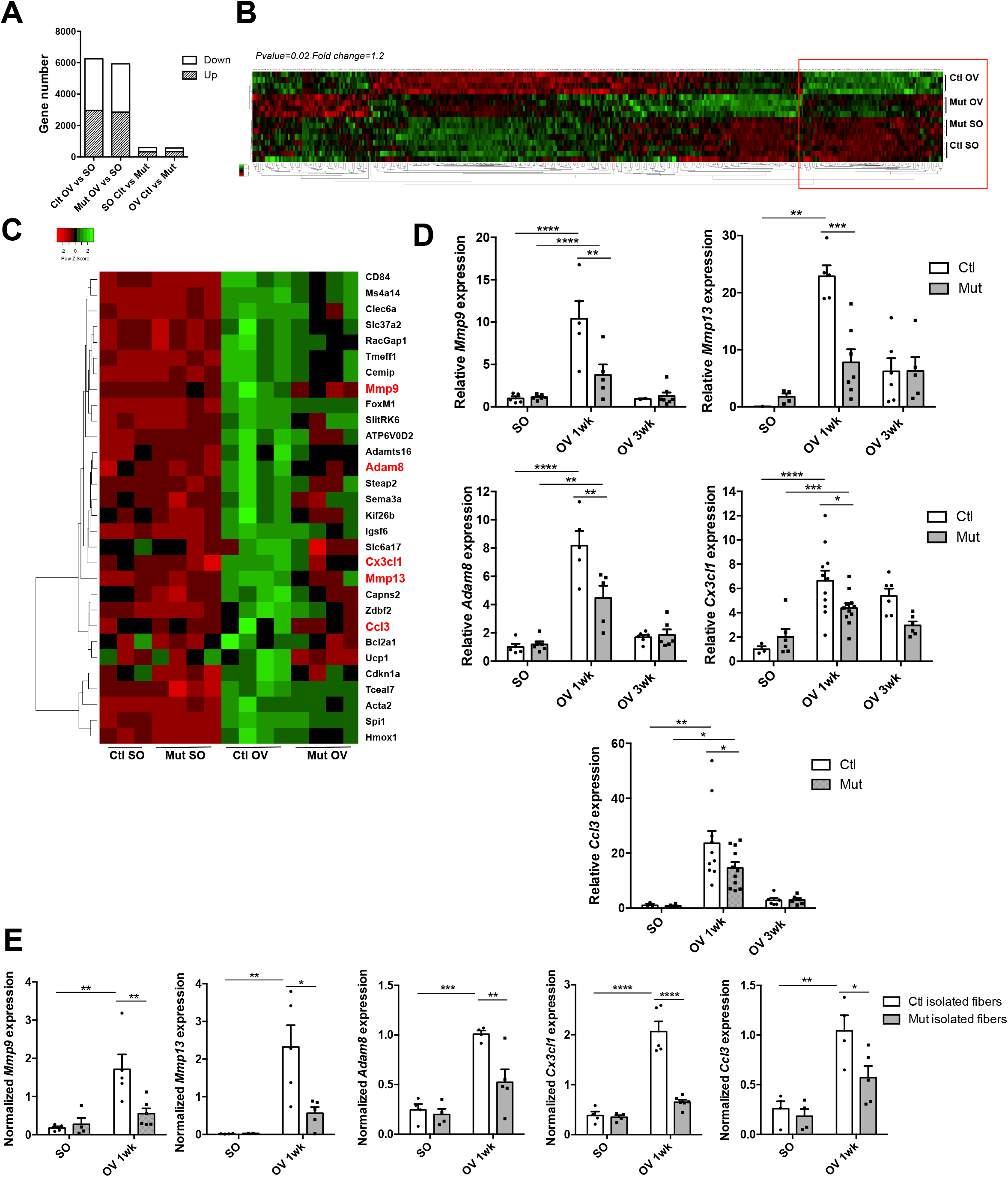
Transcriptomic analysis of control and RhoA mutant overloaded *plantaris* muscles. **(A)** Affymetrix analysis has been performed from RNA extracted from Ctl and Mut *plantaris* before (SO) and after 1wk OV. Number of genes differentially expressed (fold change>1.2; pvalue<0.05) depending on the condition SO/OV and on the RhoA expression Ctl/Mut. **(B)** Heat map representing up- and down-regulated genes between Ctl and Mut *plantaris* before and after 1wk OV. Genes whose expressions increased upon OV in Ctl are highlighted with a red box. **(C)** Close-up on the top 30 differentially expressed genes between Ctl OV and Mut OV. **(D)** Analysis of *Mmp9*, *Mmp13*, *Adam8*, *Ccl3* and *Cx3Cl1* mRNA expression by RT-qPCR in Ctl and Mut *plantaris* before (SO) and after 1 and 3wk OV (n=3-10). Data were normalized by *Hmbs* expression. **(E**) Analysis of *Mmp9*, *Mmp13*, *Adam8*, *Ccl3* and *Cx3Cl1* mRNA expression by RT-qPCR in Ctl and Mut isolated myofibers from *plantaris* muscles before (SO) and after 1 wk OV (n=4-6). Data were normalized by *Hmbs* expression. Data and mean±SEM. *pvalue<0.05, **pvalue<0.01, ***pvalue<0.001, ****pvalue<0.0001.

**Figure 4.**
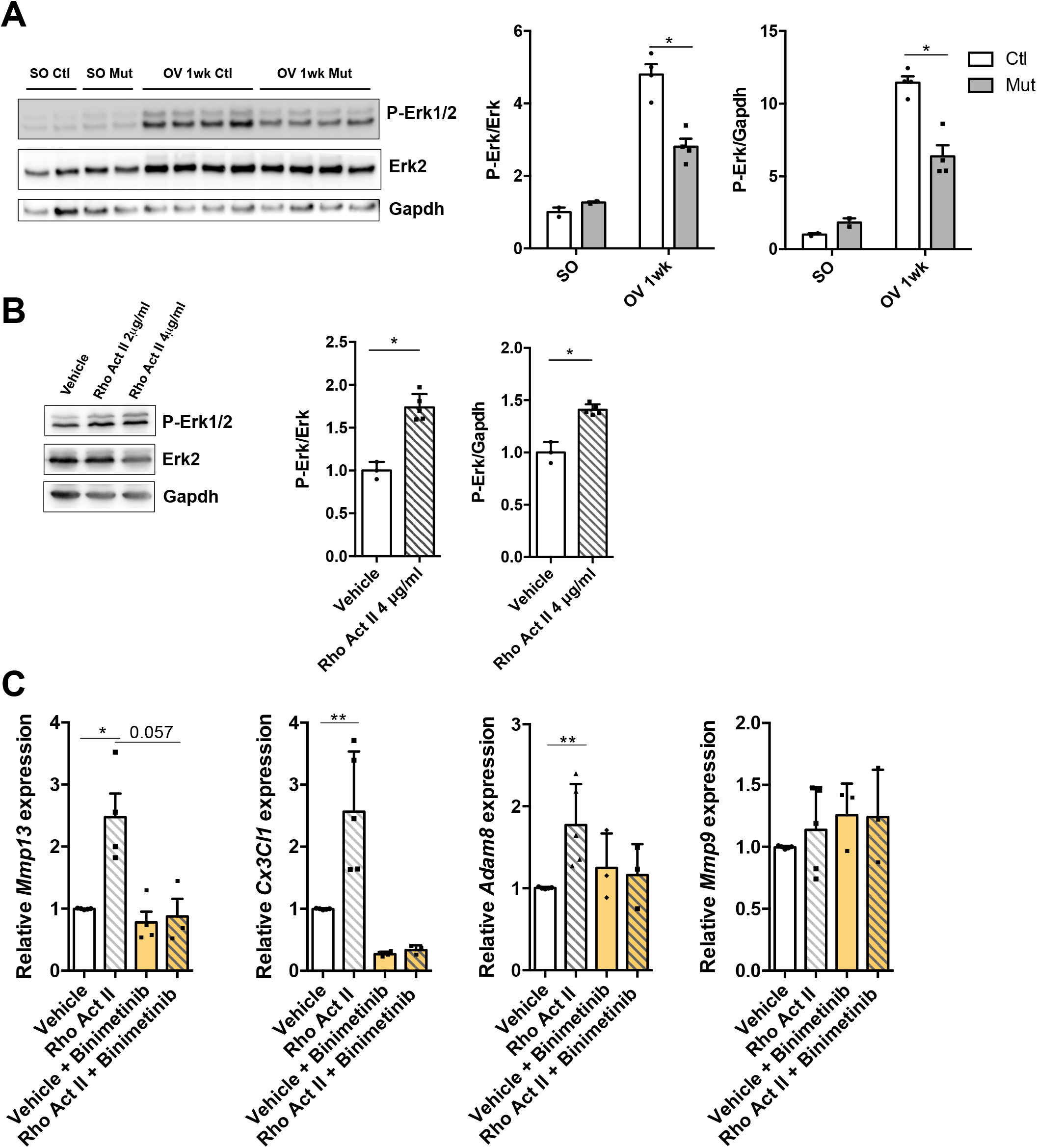
RhoA loss within myofibers impairs Erk1/2 activation upon overload-induced hypertrophy. **(A)** Phosphorylated Erk1/2 and total Erk2 were analysed by Western blot in Ctl and Mut *plantaris* before (SO) and after 1wk OV (n=4). Gapdh was used as a loading control. Ratio of the quantification of P-Erk1/2 to Erk1/2 relative to Ctl SO and ratio of the quantification of P-Erk1/2 to Gapdh are shown in the right panels. **(B)** Phosphorylated Erk1/2 and total Erk2 were analysed by Western blot in Ctl myotubes treated with Rho Activator II or vehicle for 3hr (n=3-5). Gapdh was used as a loading control. Ratio of the quantification of P-Erk1/2 to Erk1/2 and ratio of the quantification of P-Erk1/2 to Gapdh are shown in the right panels. **(C)** Analysis of *Mmp9*, *Mmp13, Adam8* and *Cx3Cl1* mRNA expression by RT-qPCR in Ctl myotubes treated with Rho Activator II (4μg/ml) and/or Erk inhibitor (Binimetenib) for 8hr (n=3-4). Data were normalized by *Hmbs* expression. Data and mean±SEM. *pvalue<0.05, **pvalue <0.01.

To investigate whether the altered expression of these genes could be linked to a lack of RhoA in myofibers or from other cell types expressing RhoA present in the muscle, we isolated single myofibers from Ctl and Mut *plantaris* before and following OV. In line with the data on bulk RNA, we showed that expression levels of *Mmp9, Mmp13, Adam8, Cx3Cl1* and *Ccl3* genes were strongly increased in Ctl single myofibers following OV and were significantly diminished in Mut (Figure 3E).

Moreover, we performed a transcriptomic analysis of gene differentially expressed between *RhoA^flox/flox^* myotubes expressing (Ad-mCherry transduced) or not RhoA (Ad-Cre-mCherry transduced). Strikingly, there was a very little overlap (41 genes) between the genes differentially expressed in myotubes in culture expressing or not RhoA and the genes differentially expressed in Ctl and RhoA Mut OV *plantaris* muscles (Mut OV 1wk vs Ctl OV 1wk). The expression of only 25 genes varied in the same direction (up or down) (Figure S3E, Table S2). *Mmp9, Mmp13, Adam8 and Cx3Cl1* genes were not among them, indicating that their expression was not affected by *RhoA* deletion in basal culture conditions (Figure S3B, Table S2). Together, these findings further bring out that the absence of RhoA in myotubes *in vitro* cannot recapitulate the *in vivo* hypertrophic setting. One parameter that is lacking *in vitro*, as compared to OV *in vivo*, is the absence of mechanical load applied to the myotubes that could be associated with low RhoA activity. Indeed it has been shown that stretch can activate RhoA in myotubes (Zhang et al., 2007). In order to activate RhoA in standard static culture conditions, we treated myotubes with a RhoA activator (Rho Activator II) and we monitored gene expression after 8hr. Importantly the expression levels of *Mmp13, Adam8*, and *Cx3Cl1* were significantly increased following RhoA activation (Figure 4C). These observations are in line with the data obtained in isolated single myofibers (Figure 3E). They suggested that at least a part of the impaired expression of these genes *in vivo* can be attributed to the specific absence of RhoA in myofibers and is thus cell autonomous. We hypothesized that the decreased expression of those genes in myofibers upon OV might participate to the incorrect hypertrophic response of mutant muscles.

### Pathways impaired by RhoA depletion in overloaded myofibers

To obtain further insights on the pathways that might be impaired by *RhoA* deletion, we focused our attention on the Upstream Regulators predicted activated or inhibited following OV in Ctl but not in Mut muscles using IPA analysis (Z-score>2 or <2, and pvalue<0.05). Several Upstream Regulators were involved in the remodeling of ECM such as Mmp9 and TnC (Table S3). Mmp9 stood as an interesting molecule because: i) it is one of the genes whose expression was the most decreased in Mut OV as compared to Ctl OV (Figure 3C) ii) it is among the Upstream Regulator predicted activated only in the Ctl and iii) it is connected to the predicted activation/inhibition of other several Upstream Regulators such as Mmps substrates TnC, Fgf, Tnf, Cxcl1 and SerpinA4 (Table S3). Mmps are proteolytic enzymes that hydrolyse components of the ECM participating to its remodeling and release many bioactive cytokines and growth factors entrapped in ECM (Page-McCaw et al., 2007). For instance, the release and activation of Tnfα and the matrix-associated growth factor Fgf by Mmp9 have been reported and consistently their activities were predicted to be activated in our IPA analysis (Table S3) (Allen et al., 2003). In the case of SerpinA4, its cleavage by Mmp9 inactivates Serpin and accordingly its activity was predicted to be decreased (Z-score = −2.17) (Table S3) (Liu et al., 2000).

In addition, several Upstream Regulators, predicted activated only in Ctl but not in Mut after OV, were associated with inflammation such as IL3, IL17RA and Gm-csf (implicated in the functional activation and survival of macrophages) and chemokines Ccl2, Cxcl1, chemokine receptors CcR1 (Ccl3 receptor) and CcR2 (Ccl2 receptor) participating to macrophage recruitment (Table S3).

Finally, several signaling pathways including Erk1 (Mapk3), Jnk (Mapk8), Stat3, Nfkb (RelA and Sn50 peptide) were predicted activated in Ctl OV *plantaris* and not in Mut OV(Table S3). We thus checked by Western blot their activity (Erk1/2, Stat3, JNK) in OV Ctl and OV Mut muscles. A difference between Ctl and Mut was only observed for Erk1/2 activity. Indeed, the increase of P-Erk1/2 following OV is significantly less in Mut as compared to Ctl muscles (Figure 4A). In order to investigate whether RhoA can activate directly Erk1/2 within myofibers, we treated Ctl myotubes *in vitro* with a Rho activator. Interestingly, a short activation of Rho GTPases (3hr) was sufficient to induce an increase of P-Erk1/2 in a dose dependent manner (Figure 4B) suggesting that the decrease of P-Erk1/2 observed in the OV Mut muscles could be partially attributed to the lack of RhoA in myofibers. Interestingly, the inhibition of Erk signaling using Binimetinib blunted the increase of *Cxcl3* and *Mmp13* expression induced by RhoA activation in Ctl myotubes (Figure 4C). Altogether, these data suggest that, upon hypertrophy, RhoA in the myofibers might affect pathways involving Erk1/2 signaling, ECM remodeling and macrophage recruitment/activity.

### RhoA in myofiber impairs ECM organization and Mmps activity

Based on the transcriptomic and IPA analyses, we hypothesized that ECM remodeling and Mmp activity might be affected by RhoA loss in the growing myofibers. We demonstrated that the global collagen content (quantified by Picro Sirius Red staining) increased in a similar extend upon OV in both Ctl and Mut muscles (Figure 5A). To further analyse ECM organization, we investigated ECM biophysical aspect by performing second harmonic generation (SHG) imaging that allows the visualization of fibrillar collagen (Figure 5B). We observed that fibrillar collagen appears more tortuous in Mut compared to Ctl 3wk after OV, as shown by the decrease of the ratio of fibrillar collagen Feret’s diameter to length (Figure 5C). Moreover it accumulated between the myofibers rather than surrounding them. These data suggest that in hypertrophic conditions RhoA impacts ECM organization in terms of tortuosity and assembly.

**Figure 5.**
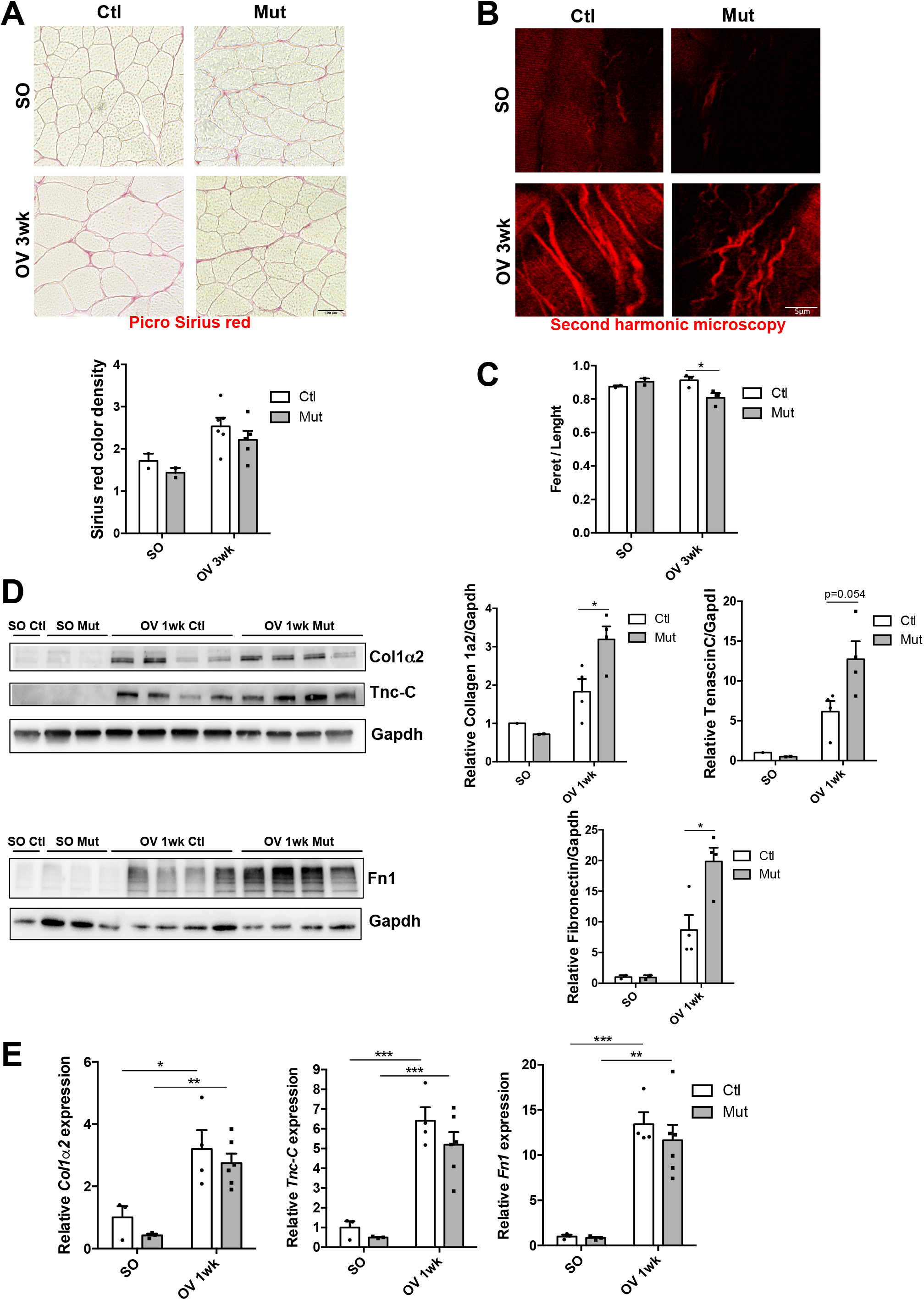
RhoA loss within myofibers impairs ECM organization upon overload-induced hypertrophy. **(A)** Picro sirius red staining of *plantaris* muscle sections from Ctl and Mut mice before (SO) and after 3wk OV. Quantification of color density was performed using CaseViewer (n=2-5). **(B)** Visualization of fibrillar collagen using second harmonic generation imaging on thick *plantaris* muscle sections from Ctl and Mut before (SO) and after 3wk OV. **(C)** Tortuosity quantification (ratio Feret to lenght of one collagen fiber) from Ctl and Mut before (SO) and after 3wk OV (n=3). **(D)** Collagen1α2, Tenascin-C and Fibronectin were analysed by Western blot in Ctl and Mut *plantaris* before (SO) and after 1wk OV. Gapdh was used as a loading control (n=3-4). Ratio of the quantification of Col1α2, TnC and Fn1 to Gapdh and relative to Ctl SO is shown in the right panel. **(E)** Analysis of *Col1α1*, *Tnc*, and *Fn1* mRNA expression by RT-qPCR in Ctl and Mut *plantaris* before (SO) and after 1 and 3wk OV (n=3-4). Data were normalized by *Hmbs* expression. Data and mean±SEM. *pvalue<0.05, **pvalue<0.01, ***pvalue<0.001, ****pvalue<0.0001.

As Mmps are key proteolytic enzymes that hydrolyze components of the ECM, we next quantified the protein amount of ECM substrates of Mmp9 and Mmp13 such as Collagen1α2 (Col1α2), TnC and Fn1 by Western blot (Figure 5D). We observed that Col1α2, TnC and Fn1 protein amounts augmented with OV in both Ctl and Mut muscles and that there was a further significant accumulation of these proteins in Mut OV muscles. Interestingly, the sharp increase upon OV in both genotypes was also seen at mRNA level indicating that this may mainly be due to increased transcription of these genes (Figure 5E). Importantly, there was no difference in transcript level between OV Ctl and OV Mut *plantaris* suggesting that the increase in Col1α2, TnC and Fn1 proteins amounts observed in the Mut might be due to their decreased cleavage by Mmp9 and Mmp13 (Figure 5E).

Thus, we could speculate that in absence of RhoA, Mmp expression diminution might affect the correct ECM protein hydrolysis and that a wrong ECM rearrangement could impair muscle growth.

### Mmps are functionally involved in compensatory hypertrophy

In order to get insights on the functional implication of Mmps in the hypertrophic process, we treated mice using the broad spectrum Mmp inhibitor GM6001 (targeting Mmp1, 2, 9, 8, 3) and we assessed its effect on compensatory hypertrophy (Figure 6A). One week post-OV, Fn1 protein amount, a common substrate of several Mmps, increased in GM6001-treated muscles, showing the *in vivo* efficacy of the drug (Figure 6B). *Plantaris* muscles of GM6001-treated mice were less hypertrophied 3wk after OV than muscles of vehicle-injected mice as assessed by their decreased weight and CSA (Figure 6C and 6D). Importantly, this altered growth was accompanied by a reduction of the number of myonuclei per myofiber (Figure 6E). Altogether these results show the important role of Mmps to achieve optimal hypertrophy in response to increased load. As Mmp inhibition phenocopied the altered growth and SC fusion observed in RhoA mutant muscles and as *Mmp9/13* expression was decreased in RhoA mutants upon OV, our findings suggest that the blunted hypertrophy of *RhoA* deleted muscles may be partially due to a decreased Mmp activity.

**Figure 6.**
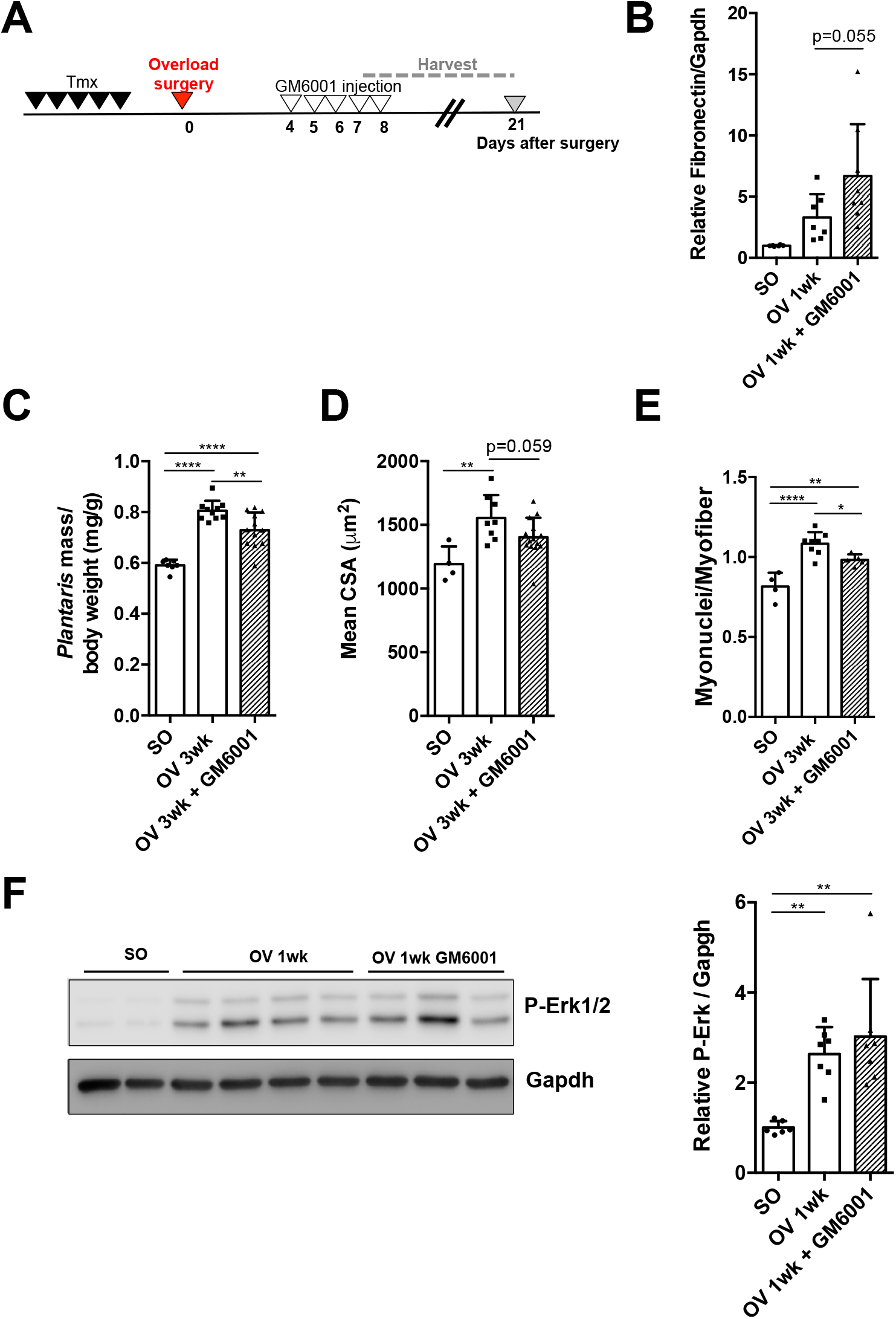
Mmps activity are important for overload-induced hypertrophy. **(A)** Ctl mice were submitted to OV and injected with Mmp inhibitor (GM6001) or vehicle from day 4 to 8 post surgery. *Plantaris* muscles were isolated after 1 and 3wk OV. **(B)** Fibronectin was analysed by Western blot in Ctl and Mut *plantaris* before (SO) and after 1wk OV. Gapdh was used as a loading control. Ratio of the quantification of Fn1 to Gapdh relative to Ctl SO is shown (n=6-8). **(C)** Ratio of *plantaris* mass (mg) to body weight (g) before (SO) and after 3wk OV in Ctl treated or not with GM6001 (n=7-14). **(D)** Mean CSA (μm^2^) before (SO) and after 3wk OV in Ctl treated or not with GM6001 (n=4-14). **(E)** Number of myonuclei within sarcolemma per myofiber in *plantaris* sections before (SO) and after 3wk OV from Ctl treated or not with GM6001 (n=4-8). Phosphorylated Erk1/2 was analysed by Western blot in Ctl treated or not with GM6001 before and 1wk OV. Gapdh was used as a loading control. Ratio of the quantification of P-Erk1/2 to Gapdh is shown in the right panel (n=3-8). Data and mean± SEM. *pvalue<0.05, **pvalue<0.01, ****pvalue<0.0001.

Mmp processing can release ECM-tethered growth factors and bioactive cytokines that can indirectly activate different signaling pathways. Therefore, we wondered whether the decreased activation of Erk1/2 observed in RhoA mutant OV muscles (Figure 4A) could be partially attributed to the diminished Mmp expression/activity. Importantly, GM6001- and vehicle-treated muscles displayed similar levels of P-Erk1/2 upon OV suggesting that the alteration of Erk signaling is not downstream of the Mmps (Figure 6F).

### RhoA in overloaded myofibers impacts macrophage recruitment implicated in muscle growth

In absence of RhoA, we showed a decreased expression of *Ccl3* and *Cx3cl1* by the growing myofibers (Figure 3D and 3E). These two chemokines/chemo-attractants are implicated in inflammation by recruiting and activating macrophages (Zhao et al., 2016; Maurer et al., 2004). In addition, IPA analysis highlighted several Upstream Regulators activated in OV Ctl linked to the inflammatory response (Table S3). Therefore, we investigated the number of macrophages (F4/80^+^) present in muscle sections following compensatory hypertrophy. One week after OV, there was a 10-fold increase in macrophage number in Ctl muscles that came back almost to SO levels 3wk post OV, showing a transient and strong inflammatory response in the muscle following increased workload (Figure 7A). Strikingly, macrophage number was significantly reduced in Mut compared to Ctl 1wk post OV (Figure 7A). This prompted us to hypothesize that RhoA in the myofibers may influence macrophage recruitment (by controlling indirectly *Ccl3* and *Cx3cl1* expression levels) in the muscle tissue following mechanical load increase.

**Figure 7.**
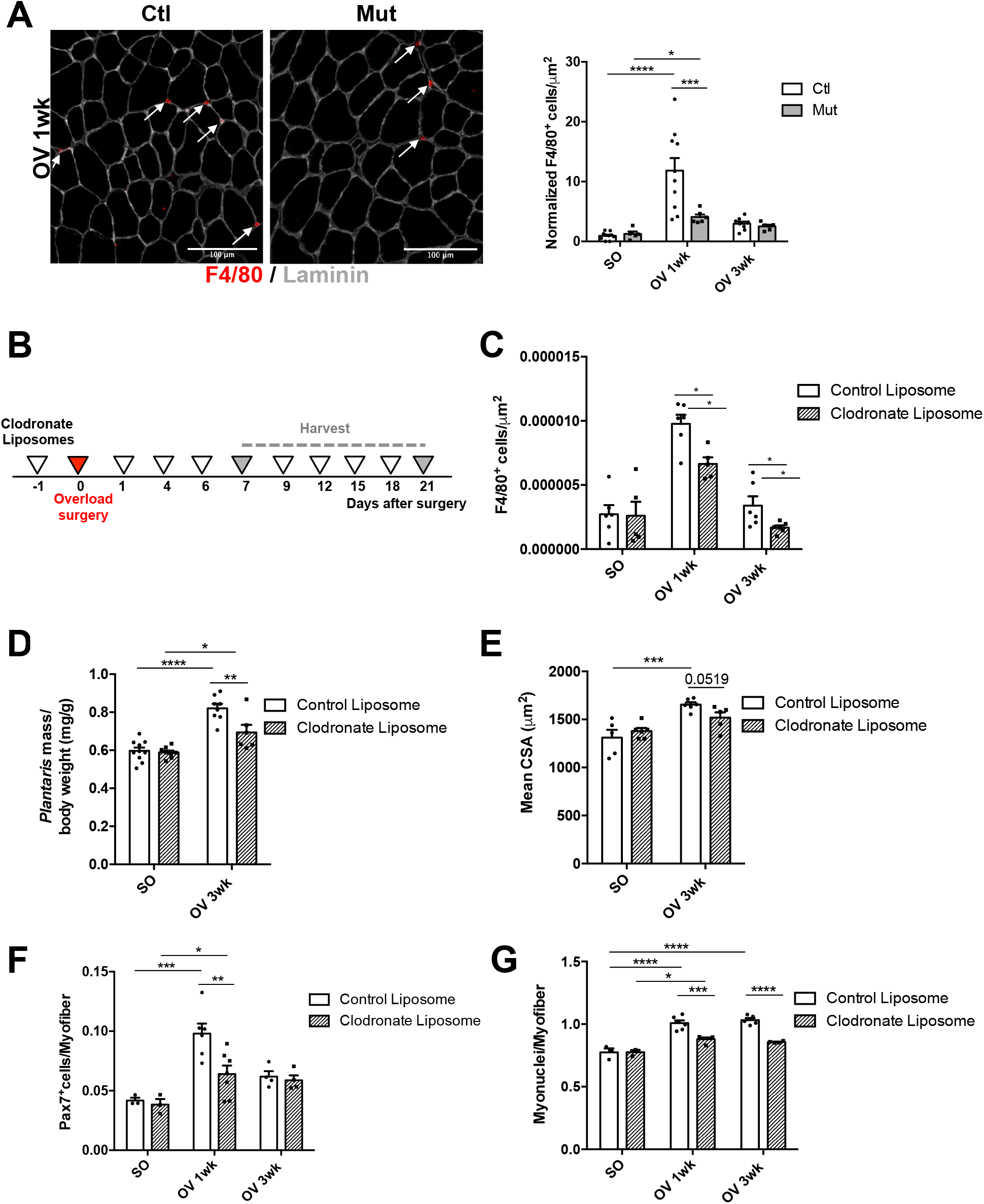
RhoA loss whithin myofiber impairs macrophage recruitment. **(A)** *Plantaris* sections were immunostained for F4/80 (red), Laminin (grey) and nuclear staining with DAPI for Ctl and Mut after 1wk OV. White arrows indicated F4/80^+^/ DAPI macrophages. Number of macrophages (F4/80^+^) per μm2 before (SO) and after 1 and 3wk OV relative to Ctl SO are quantified in the right panel (n=5-6). **(B)** Ctl mice were injected with Clodronate-loaded liposomes or control liposomes one day before OV and at day 1, 4, 6, 9,12,15,18 post OV. *Plantaris* muscles were harvested after 1 and 3wk OV. **(C)** Number of macrophages (F4/80^+^) per μm^2^ before (SO) and after 1 and 3wk OV in *plantaris* sections from Ctl treated with Clodronate loaded liposomes or control liposomes (n=5-6). **(D)** Ratio of *plantaris* mass (mg) to body weight (g) before (SO) and after 3wk OV in Ctl treated with Clodronate loaded liposomes or control liposomes (n=6-10). **(E)** Mean CSA (μm^2^) before (SO) and after 3wk OV in Ctl treated Clodronate loaded liposomes or control liposomes (n=5-6). **(F)** Number of Pax7^+^ cells per myofiber in *plantaris* sections before (SO) and after 1 and 3wk OV from Ctl treated with Clodronate loaded liposomes or control liposomes (n=3-7). **(G)** Number of myonuclei within sarcolemma per myofiber in *plantaris* sections before (SO) and after 1 and 3wk OV from Ctl treated treated with Clodronate loaded liposomes or control liposomes (n=4-8). Data and mean±SEM. *pvalue<0.05, **pvalue<0.01, **pvalue<0.01, ***pvalue<0.001, ****pvalue<0.0001.

It has been shown that macrophages are very important to ensure the proper regeneration of muscle by controlling SC behaviours such as proliferation and fusion (Chazaud, 2020). To investigate whether macrophages play a functional role during hypertrophy, we depleted the macrophages by injecting intraperitoneally Clodronate liposomes that induce macrophage death (Figure 7B). First, we checked macrophage depletion efficacy by counting the number of macrophages on *plantaris* sections 1wk post OV and showed a 30% and a 50% decrease 1wk and 3wk post OV, respectively (Figure 7C). This partial depletion was sufficient to blunt hypertrophic growth of *plantaris* muscles 3wk after OV as evidenced by the reduced *plantaris* weight and CSA in macrophage-depleted mice (Figure 7D, 7E). Moreover macrophage depletion led to decreased number and fusion of SCs (Figure 7F, 7G), strongly suggesting that, in hypertrophic conditions, macrophages modulate SC behaviour. These results are reminiscent of those observed in RhoA Mut overloaded muscles, in which SC behaviours are altered (Figure 2), and suggest that RhoA within the myofiber may partially control muscle growth by favouring the recruitment of macrophages in overloaded muscles.

## Discussion

Taking advantage of a genetic model that allows the deletion of *RhoA* in myofibers and not in SCs, we demonstrated that RhoA is needed for skeletal muscle hypertrophy and for the recruitment of SCs to the growing fibers. We provided evidence for the implication of RhoA within myofibers in the building of a permissive microenvironment for muscle growth and for SC accretion through ECM remodeling and macrophage recruitment. At the molecular level, we propose that, in response to increased workload, RhoA controls in a cell autonomous manner Erk1/2 activation and the expression of ECM regulators such as *Mmp9/Mmp13/Adam8* and of macrophage chemo-attractants such as *Ccl3*/*Cx3cl1* (Figure S4).

Hypertrophic growth of skeletal muscle relies on both the increase of net protein content and the addition of new nuclei in the growing myofibers provided by SCs. We showed that, in absence of RhoA, the hypertrophy defect is not accompanied by a deregulation of protein synthesis/degradation balance or by a compromised Akt signaling, but mostly by a deficiency of SC fusion. We propose that, upon OV-induced hypertrophy, the altered recruitment of SC to *RhoA* mutant myofibers may be responsible for their impaired growth. Indeed, even if the role of SCs in skeletal muscle hypertrophic growth has been debated, the necessity of competent fusing SCs to have a correct muscle hypertrophy has been clearly demonstrated using genetic mouse models in which SCs are unable to fuse. For instance, SC deletion of *Myomaker*, a muscle-specific membrane protein necessary for fusion of muscle progenitors, compromised muscle hypertrophic response (Goh & Millay, 2017). In addition, loss of Srf in SCs, a master regulator of F-actin scaffold, impaired OV-induced muscle growth by the absence of SC fusion (Randrianarison-Huetz et al., 2018).

RhoA is an important small GTPase protein regulating contractility, actin cytoskeleton rearrangement, and actin polymerization through its downstream effector Rho-associated coiled-coil protein kinase and mammalian Diaphanous protein among many others. Moreover RhoA activation responds to mechanical stimuli and mechanical force transmitted and mediated by cell-matrix and cell-cell-interaction (Lessey et al., 2012). In cultured C2C12 myotubes, it has been shown that cyclic mechanical stretch increased RhoA activity (Zhang et al., 2007) suggesting that RhoA participates to the mechanical stretch response in myotubes. To identify the molecular mechanisms underlying the lack of hypertrophic growth of *RhoA* mutant myofibers, we compared the data of microarray experiments performed *in vivo* on OV *plantaris* Ctl and Mut muscles and the data of microarrays performed on differentiated myotubes expressing or not RhoA. Surprisingly, the intersection set was very small, suggesting that the pathways regulated by RhoA during OV *in vivo* are very different from those operating in myotubes cultured in standard static conditions. None of the following genes *Mmp9, Mmp13, Adam8, Ccl3, Cx3Cl1*, which were differentially expressed between Clt and Mut overloaded whole *plantaris* muscles and isolated myofibers, was expressed differentially in myotubes expressing or not RhoA. Additionally, we were not able to reproduce the fusion defects of Ctl myocytes with RhoA mutant myotubes *in vitro*. These discrepancies between *in vivo* and *in vitro* settings highlight the need of mechanical inputs to trigger RhoA-dependent muscle hypertrophy. In addition, in response to increased load, RhoA in the myofibers may affect other cell types present in the muscle (FAPs, macrophages), but absent *in vitro*, that may participate to muscle growth. Altogether, this emphasizes the importance of *in vivo* models to analyse RhoA contribution to muscle hypertrophy.

It has been shown that Rho GTPases, and in particular RhoA, are potent activators of Srf transcriptional activity (Sotiropoulos et al., 1999). In addition, we have previously shown that Srf within the myofibers is required for muscle OV-induced hypertrophy by exerting a paracrine control on SC recruitment to the growing fibers through Ptgs2/IL4 (Guerci et al., 2012). We could wonder whether a decrease of Srf activity could be responsible for the impaired hypertrophic growth of *RhoA* deleted muscles. Several data argue against this possibility. First, our *in vivo* and *in vitro* transcriptomic data on muscles upon OV and on myotubes suggested that RhoA governs downstream signaling pathways independently from Srf, as almost none of the *bona fide* Srf target genes are down-regulated when RhoA is absent. For instance, *Ptgs2* (a Srf target gene), which expression increases during OV in Ctl muscles and is blunted in Srf mutants, displays a similar expression in Ctl and RhoA Mut *plantaris* 1wk post OV. In cultured myotubes, *RhoA* deletion did not affect the expression of Srf target genes such as *Egr1*, *Myl9, Cnn2, Ptgs2* and *Acta1* (Guerci et al., 2012; Randrianarison-Huetz et al., 2018). Second, the expression of several “RhoA responsive” genes such as *Mmp13, Adam8* and *Cx3Cl1* that increased following chemical activation of RhoA *in vitro* on cultured myotubes and that decreased in overloaded RhoA Mut muscles, remained unaffected by constitutive Srf activation in myotubes (induced by Ad-SrfVP16 transduction) (Guerci et al., 2012). Thus, we propose that the impeded muscle growth of RhoA mutant is likely not due to a decreased Srf activity and that Srf-independent signaling pathways are operating downstream RhoA upon hypertrophy.

In the present study, we showed that genes involved in the ECM remodeling, like *Mmp9* and *Mmp13* are among the genes mostly down-regulated in overloaded muscles in absence of RhoA. In addition, *in vivo* Mmps inhibition by a broad-spectrum Mmp inhibitor impaired OV-induced hypertrophy and SC accretion, and phenocopied RhoA mutants. Interestingly, Mmp inhibitor GM6001 was shown to alter SC number following muscle laceration-induced injury (Bellayr et al., 2013). Thus, we propose that *Mmp9* and *Mmp13* decreased expression may contribute to the altered muscle growth of RhoA mutants by modulating SC fusion. Many studies were conducted to show the importance of Mmps on muscle growth. Fiber size is decreased in adult hindlimb muscles from *Mmp9* null mice (Mehan et al., 2011) and increased in muscles harbouring a specific overexpression of *Mmp9* (Dahiya et al., 2011). In the context of hypertrophic growth, Peterson laboratory recently proposed that Mmp9 levels in muscle fibers should be finely regulated (through miRNA secreted by SCs among others) to have a correct ECM integrity and to facilitate the long-term hypertrophic response; too much or not enough Mmp9 being deleterious (Fry et al., 2020). In line, continuous unregulated high Mmp9 activity disrupted SC niche through ECM damages (Chenette et al., 2016). *Mmp13* null mice did not exhibit any histological or functional muscle deficits, however muscle hypertrophy caused by increased IGF-1 were impaired (Smith et al., 2020). Mmp9 and 13 have also been shown to modulate myoblast migration *in vitro* (Lei et al., 2000). It is reasonable to assume that the remodeling of ECM by Mmps enables more efficient movement of myoblasts toward myofibers and thus contributes to SC recruitment upon growth.

While searching for additional players responsible for the impaired growth of RhoA mutant muscles, we focused our interest on two potent macrophage attractants: *Cx3cl1* and *Ccl3* whose expression is increased upon OV in Ctl and not in Mut. Data obtained in Ctl and Mut isolated myofibers and in myotubes treated with a Rho activator plus an Erk inhibitor further demonstrated that RhoA stimulated the expression of these chemokines in a myofiber cell-autonomous and in an Erk1/2-dependent manner. We do not know whether Cx3cl1 and Ccl3 influence SCs directly, but we can strongly relate their decreased expressions to a significant impairment of macrophages recruitment in RhoA mutants. Indeed, OV induced a transient accumulation of macrophages in Ctl *plantaris* muscles that was blunted in Mut. In human, it has been shown that resistance exercise increases the expression of chemotaxic factors including *Cx3cl1*, mobilizes monocytes (Della Gatta et al., 2014; Jajtner et al., 2018; Strömberg et al., 2016) and that macrophage number positively correlates with myofibers hypertrophy (Walton et al., 2019). Knowing the multifaceted roles of macrophages during muscle regeneration to support tissue recovery, in particular by promoting SC differentiation (Chazaud, 2020), we proposed that macrophages may as well participate to muscle hypertrophy by facilitating SC recruitment to myofibers. By depleting macrophages in Ctl mice, we showed that macrophages were required for muscle hypertrophy. Our results are in agreement with those of DiPasquale who showed that macrophages depletion also negatively affected muscle hypertrophic growth (DiPasquale et al., 2007). However we provide additional data concerning the positive action of macrophages on SC number and fusion.

It is important to note that the absence of RhoA in the myofibers is sufficient to affect the behaviour of RhoA expressing SCs, highlighting the importance of RhoA signaling in myofibers for the remodeling of SC microenvironment upon hypertrophy. How the SC niche is remodeled during hypertrophy process remains poorly examined. It was already proved that, after increased load, myofibers secrete myokines like IL-4 and IL-6 that are able to influence SC proliferation and fusion (Guerci et al., 2012, Serrano et al, 2008, Horsley et al, 2003). In addition, important transcriptional modifications of the expression of several genes encoding ECM components (such as *Collagens*, *Fn1* and *TnC*) have been reported shortly after OV, in this study (1wk) and Carson ‘s study (3 days post OV in rat) (Carson et al., 2002). By affecting the expression of *Mmp9* and *Mmp13*, RhoA might promote an important global ECM rearrangement and a modification of SC niche upon OV that could influence SC behaviour such as fusion. We show protein accumulation of several Mmp9/13 ECM protein substrates such as Col1α2, TnC and Fn1. Moreover, analysis of collagen fibrils structure in OV Mut muscles displayed a significant increase in fibrillar collagen tortuosity as compared to Ctl. These data suggest that the correct rearrangement of ECM/collagen is crucial to build a SC microenvironment appropriated for muscle growth and SC fusion. Interestingly, both TnC and Fn1 have been shown to remodel and adapt SC niche (Tierney et al., 2016) and to regulate a feedback signal to promote symmetric division and replenish SC pool (Bentzinger et al., 2013), respectively. In both cases, the proteins are produced by SCs that auto-regulate their own niche. Because in our instance SCs are “wild type”, we could speculate that this auto-regulation is not sufficient alone but should be associated with enzyme activity for ECM remodeling from the myofiber. Of note, as macrophage number is decreased in RhoA Mut compared to Ctl muscles upon OV, on top of a myofibers intrinsic deficit, one part of the defective ECM reorganization occurring in Mut could be attributed to macrophages which also secrete Mmp9 in a context of muscle regeneration (Rahman et al., 2020).

In conclusion we have described an important role for RhoA within myofibers in regulating muscle hypertrophic growth and SC accretion. We propose that it is achieved through a modification of SC microenvironment by a fine ECM remodeling and by inflammatory cells recruitment. It will be important in future works to define more comprehensively the molecular pathways involved and the specific cellular contributions as unraveling the pathways involved in the control of muscle mass, in particular in response to increased mechanical load, may be essential to identify and design treatments able to alleviate loss of muscle mass during disuse or aging.

## Acknowledgments

We thank the Cybio, Genomi’IC and Animal care core facilities of Cochin Institute (Paris, France). We are grateful to Evelyne Bloch-Galego, Florian Britto for critical reading of the manuscript and So-ichiro Fukada for technical expertise. SHG images acquisition and analysis were performed at the IMAG’IC Facility, supported by the National Infrastructure France BioImaging (grant ANR-10-INBS-04) and IBISA consortium.

Financial support to this work was provided by the AFM (n°22365 and 22776), IdEx Emergence Université de Paris (IP 2019-030) the Agence Nationale pour la Recherche (Myolinc, ANR-R17062KK), the Institut National de la Santé et de la Recherche Médicale (INSERM) and the Centre National de la Recherche Scientifique (CNRS).

## Author contributions

CN, KK, designed and carried out experiments, analyzed results and wrote the manuscript. LD, FJ, TG conducted and analyzed experiments. PM provided expertise and financial support. VR provided expertise and wrote the manuscript. AS designed and carried out experiments, analyzed results, wrote the manuscript and provided financial support.

## STAR Methods

### Mouse protocols

*RhoA^flox/flox^* mice are homozygous for RhoA floxed alleles harboring LoxP sites flanking exon 3 of endogenous *RhoA* gene (Lauriol et al., 2014). HSA-*Cre-ER^T2^* transgenic mice express Cre-ERT2 recombinase under the control of human skeletal actin promoter (Schuler et al., 2005). Tg:Pax7-nGFP transgenic mice express nuclear localized EGFP under the Pax7 promoter (Sambasivan et al., 2009).

To investigate the effect of myofiber-specific RhoA-deletion in adult muscle, the mouse strain following mice were generated: HSA-*Cre-ER^T2^*:*RhoA^flox/flox^.* In all experiments, 3-month-old HSA-*Cre-ER^T2^*:*RhoA^flox/flox^* mice were given five intraperitoneal (i.p.) tamoxifen (Tmx, 1mg/day; MP Biomedicals) injections to induce *RhoA* deletion and were referred as mutant mice (Mut). *RhoA^flox/flox^* mice injected with Tmx were used as control mice (Ctl).

No statistical differences in body weights were observed after Tmx in HSA-*Cre-ER^T2^*:*RhoA^flox/flox^* and *RhoA^flox/flox^* mice up to 2 months post Tmx injection.

Mice were genotyped by PCR using the following primers: Cre-F: 5’-CCTGGAAAATGCTTCTGTCCG-3’; Cre-R: 5’-CAGGGTGTTATAAGCAATCCC-3’; RhoAlox-F: 5’-AGG GTT TCT CTG TAC GGT AGTC-3’; RhoAlox-R: 5’-GCA GCT AGT CTA ACC CAC TACA-3’. Overload-induced hypertrophy (OV) of *plantaris* muscles of control (Ctl) and mutant (Mut) mice was induced through the incapacitation of *soleus* and *gastrocnemius* muscles by sectioning their tendon. This procedure was achieved in both legs. During the process of OV, Mut mice were injected with TMX at day 3, 5 and 7 post OV. At the indicated time (1 and 3wk post OV), *plantaris* muscles were dissected and subsequently processed for histological analyses. When indicated mice were administered intraperitoneally with 50 or 20 μg/g EdU (Life Technologies) or with 20μg/g GM6001 (Merck Millipore).

All animal experiments were conducted in accordance with the European guidelines for the care and use of laboratory animals and were approved by the institutional ethic committee and French Ministry of Research (number A751402).

### Macrophage depletion

Clodronate liposome and control liposome (Clophosome-A and control liposome anionic) were purchased from ChemQuest. Mice were injected with Clodronate- or control liposomes (0.15 ml administrated intraperitoneally i.p.) 1 day prior overload-induced hypertrophy. Additionally, liposomes (Clodronate- or control) were continued to be administrated day 1, 4, 6, 9, 12, 15 and 18 post-surgery (0.1ml, i.p.). The muscles were harvested 7 days or 21 days post compensatory hypertrophy.

### SUnSET

The SUnSET, or SUrface SEnsing of Translation, technique allows estimation of global protein synthesis in tissue of living animals, specifically involves the use of an anti-puromycin antibody for the immunological detection of puromycin-labelled peptides. Briefly we injected mice with puromycin (Sigma) at 0.04μg/g body mass via an i.p. injection 30min before samples harvesting. Western blot was then performed with a mouse anti-puromycin antibody (1/1000 Merck Millipore) and HRP-conjugated anti-mouse secondary antibody. Puromycin incorporation was measured by Fusion FX system (Vilber company) with Comassie blue staining as loading control.

### Multiphoton imaging and characterization of ECM topography and architecture

*Plantaris* muscle was excised and placed in 2% paraformaldehyde (PFA) for 1hr at 4°C and then included in agarose low gelling 6%. Muscle was then sliced in thick sections (300 μm) using a Leica VT1200 S vibratome (Leica Microsystems, Wetzlar, Germany). Leica SP5 multiphoton microscope (Leica Microsystems, Wetzlar, Germany) coupled with an Ultra II Chameleon Coherent TI:sapphire laser (Coherent, Saclay, France) was used to image collagen fibers by second harmonic generation microscopy. The laser beam was circularly polarized and a Leica Microsystems HCX IRAPO 25x/0.95 W objective was used. From z-stack images, the tortuosity was calculated using collagen fibrils Feret’s diameter divided by length (ImageJ software) of at least 50 single collagen fibrils per sample, as described in (Stearns-Reider et al., 2017)(Stearns-Reider et al., 2017).

### Primary muscle cell culture and recombinant virus transduction

Primary cultures were derived from hindlimb muscles of *RhoA^flox/flox^ mice* harboring Pax7*-nGFP* transgene that allowed the prospective selection by flow cytometry (Fluorescence Activated Cell Sorting or FACS) of satellite cells as described in (Randrianarison-Huetz et al., 2018). In standard conditions, myoblasts were grown in growth medium (DMEM/F12, 2% Ultroser G (PALL Life Sciences), 20% Fetal Calf Serum) on plastic dishes coated with 0.02% Gelatin. For differentiation, myoblasts were seeded in Matrigel-coated dishes and cultured in differentiation medium (DMEM/F12, 2% Horse Serum). When indicated cells were treated with Rho activator II CN03 (Cytoskeleton, 2 and 4μg/ml) and/or Erk inhibitor Binimetenib (Selleckchem, 6μM).

To induce in culture the excision of the floxed *RhoA* allele in myotubes, *RhoA^flox/flox^* myoblasts were differentiated for 3 days and then transduced with Ad-mCherry or Ad-Cre-mCherry adenoviruses (100MOI, Vector Biolabs).

To generate a stable *RhoA^flox/flox^* cell line expressing GFP, RhoA*^flox/flox^* myoblasts were transduced twice with recombinant Lenti-GFP (100MOI, Vector Biolabs) and 8μg/ml of polybrene then selected by FACS sorting.

### Cell mixing fusion assays

To analyze fusion, myotubes (at day 2 of differentiation) were transduced with Ad-Cre-mCherry or Ad-mCherry. Two days post transduction, they were co-cultured with Lenti-GFP transduced myocytes (myoblasts incubated overnight in DM). Two days later, cultured cells were fixed for 8min in 4% PFA and nuclei were stained using DAPI and then mounted in Dako Fluorescence Mounting Medium and kept at 4°C until image acquisition. Fusion events were scored by counting the dual-labeled cells (green cells GFP^+^/ mCherry^+^). The number of fusion events was normalized by the total number of nuclei.

### Muscle section immunostaining

*Plantaris* muscles were collected and snap-frozen in liquid nitrogen-cooled isopentane. Eight μm-thick muscle sections were fixed in 4%PFA for 8min at room temperature and blocked overnight at 4°C in PBS 1x, 10% Horse serum, 0.5% Triton X-100. Then, they were incubated with primary antibodies overnight at 4°C in PBS 1x, 10% Horse serum, 0.5% Triton X-100. The following primary antibodies were used: mouse anti-Dystrophin (Novocastra, NCL-Dys2, 1/50), anti-F4/80 (Invitrogen, MA1-91124, 1/400) or rabbit anti-Laminin (Sigma, L9393, 1/200). After washes in PBS 1x, sections were incubated with secondary antibodies for 1hr at room temperature. The following secondary antibodies used were goat anti-mouse IgG1 Alexa 488 (ThermoFisher, A21121, 1/1000) and donkey anti-rabbit Alexa 546 (LifeTechnologies, A10040, 1/1000). Nuclei staining were performed using DAPI. Muscles sections were then mounted in Dako Fluorescence Mounting Medium and kept at 4°C until image acquisition.

For Pax7 staining, muscle sections were fixed in 4%PFA for 8min at RT and permeabilized in ice-cold methanol for 6min. Muscle sections were treated with Antigen Unmaking Solution pH6 (Vector, H-3300) 15min at 95°C and cooled on ice for 30min. Blocking and incubation with primary and secondary antibodies were conducted as described in the previous paragraph. Primary mouse anti-Pax7 antibody (Santa Cruz, sc-81648) was used at dilution 1/50.

EdU detection was performed using Click-iT® EdU Alexa Fluor® 647 kit, according to the manufacture’s instructions (Life Technologies).

### Western blot analysis

*Plantaris* muscles were lysed in RIPA buffer (Sigma) and proteins were separated through denaturating SDS-PAGE electrophoresis using Mini-Protean TGX precast gels 4-15% (Biorad) and transferred on PVDF or Nitrocellulose (0.2μm, Biorad) membrane using the Trans-Blot turbo transfer system (Biorad). Membranes were blocked with 5% skinned milk in TBS-1% Tween (TBST) 1hr at room temperature and probed overnight at 4°C with primary antibodies in TBST 5% BSA. The following antibodies were used: mouse anti-RhoA (Santa Cruz, sc-418, 1/150), rabbit anti-P-Akt (Cell Signaling, #9271S, 1/1000), anti-Col1α2 (Abcam, ab34710, 1/1000) Fibronectin (Sigma F3648, 1/1000), anti-Erk2 (Cell Signaling, #9102, 1/1000), anti-P-Erk1/2 (Cell Signaling, #9101, 1/1000), rat anti-TenascinC (Invitrogen, MA1-26778, 1/200), and mouse anti-Gapdh (Cell Signaling, D16H11, 1/1000). Following washing in TBST, membranes were hybridized with goat anti-mouse and goat anti-rabbit secondary antibodies coupled to HRP (ThermoFisher, 62-6520 and A27036, 1/10 000). Proteins were revealed using SuperSignal West Femto substrate (ThermoFisher).

### Image acquisition

Fluorescence images were acquired using an Olympus BX63F microscope with 10x (UplanFL, NA 0.3) and 20x objectives (UPLSAPO, NA 0.75), coupled with an ORCA-Flash4.0 LT camera (Hamamatsu), or using a Zeiss Axiovert 200M microscope with 5x (PLANFLUAR, NA 0.25) and 20x objectives (LD PLANNEOFLUAR, NA 0.4), coupled with a CoolSnap-HQ camera (Photometrics), or using a Spinning Disk confocal microscope (Yokogawa X1) with a 100x oil-immersion objective (HCX PL APO, NA 1.47), coupled with a CoolSnap-HQ^2^ camera (Photometrics) and Metamorph 7.7.5 (Molecular Devices). Images were merged and edited in ImageJ. Background was reduced using brightness and contrast adjustments applied to the whole image.

### Morphometric analysis

Myofiber cross section area (CSA) was analyzed by using immunostaining of Dystrophin, marking myofiber sarcolemma, and then using MuscleJ tool (Mayeuf-Louchart et al., 2018). Between 600 and 800 myofibers were analyzed. For the quantification of the number of nuclei per myofibers ImageJ was used and at least 500 myofibers were counted per muscle.

### RNA extraction and RT-qPCR

Total RNA was extracted using TRIzol reagent and reverse-transcribed with SuperScript III reverse transcriptase (Invitrogen). cDNA was synthesized from 1μg of RNA. Quantitative PCR analysis was performed using a Light Cycler (Roche) according to the manufacturer’s instructions using a SYBR Green I kit (Roche). Values were normalized using *Hydroxymethylbilane synthetase* (*Hmbs*). The following primers were used: RhoA-F 5’-AACCTGTGTGTTTTCAGCACC-3’ ; RhoA-R 5’-ACCTCTGGGAACTGGTCCTT-3’ ; MuRF1-F 5’-GAATAGCATCCAGATCAGCAG-3’ ; MuRF1-R 5’-GAGAATGTGGCAGTGTTTGCA-3’ ; MAFbx-F 5’-TGTGGGTGTATCGGATGGAGA-3’ ; MAFbx-R 5’-CTGCATGATGTTCAGTTGTAAGC-3’ ; Mmp9-F 5’-TCCTACTCTGCCTGCACCACTAAAG-3’ ; Mmp9-R 5’-CTGTACCCTTGGTCTGGACAGAAAC-3’, Mmp13-F 5’-AGTTGACAGGCTCCGAGAAA-3’ ; Mmp13-R 5‘-CACATCAGGCACTCCACATC-3’ ; Adam8-F 5’-GCAGGACCATTGCCTCTACC-3’ ; Adam8-R 5’-TGGACCCAACTCGGAAAAAGC-3’ ; Ccl3-F 5’-TTCTCTGTACCATGACACTCTGC-3’. Ccl3-R 5’-CGTGGAATCTTCCGGCTGTAG-3’ ; Cx3Cl1-F 5’-ACGAAATGCGAAATCATGTGC-3’ ; Cx3Cl1-R 5’-CTGTGTCGTCTCCAGGACAA-3’ ; Col1α2F 5’-GTAACTTCGTGCCTAGCAACA-3’ ; Col1α2R 5’-CCTTTGTCAGAATACTGAGCAGC3’ ; TnC-F 5’-CACACACCGCATCAACATCC-3’ ; TnC-R 5’-GACGACTTCTGCAGCTTGGA-3’, Fn-F: 5’-GGCCACACCTACAACCAGTA-3’ ; Fn-R: 5’-TCGTCTCTGTCAGCTTGCAC-3’ ; Hmbs-F: 5’-TGCACGATCCTGAAACTCTG-3’; Hmbs-R 5’-TGCATGCTATCTGAGCCATC-3’.

### Affymetrix microarrays

Microarray analysis was performed from: i) four *plantaris* muscles coming from four different mice per group. Four conditions (groups) were analyzed: SO Ctl, 1wOV Ctl, SO Mut, 1wOV Mut; ii) three independent *RhoA^flox/flox^* differentiated myoblast cultures transduced with Ad-mCherry or Ad-Cre-mCherry adenoviruses for 2 days. Total RNAs were obtained using TRIzol reageant and DNAse treatment (Qiagen). RNA integrities were certified on bioanalyzer (Agilent). Hybridation to Mouse Gene 2.0-ST arrays (Affymetrix) and scans (GCS3000 7G expression Console software) were performed on the Genom’IC plateform (Institut Cochin, Paris). Probe data normalization and gene expression levels were processed using the Robust Multi-array Average (RMA) algorithm in expression Console software (Affymetrix). Gene ontology analysis was performed using Ingenuity (IPA) software.

### Statistical analysis

Quantitative data sets were analyzed using an unpaired non-parametric Mann Whitney test (Fig. 1B; Fig. 4A, 4B and 4C; Fig. 6B and 6F; Fig. 7E), unpaired t-test with Welsh correction (Fig. 5D; Fig. S3B), two-way ANOVA with Tukey’s multiple comparisons test (Fig. 1C, 1D, 1F and 1G; Fig. S1A, S1B and S1C; Fig. 2A, 2B, 2C, 2E and 2H; Fig. S2; Fig. 3D and 3E; Fig. 5E; Fig. 7A, 7C, 7D, 7E, 7F and 7G), two-way ANOVA with Sidak’s multiple comparisons test (Fig. 1D), one-way ANOVA with Tukey’s multiple comparisons test (Fig. 6C, 6D and 6E), using GraphPad Prism 6.0 software. Statistical significance was set at a pvalue<0.05.

---

**Figure S1.**
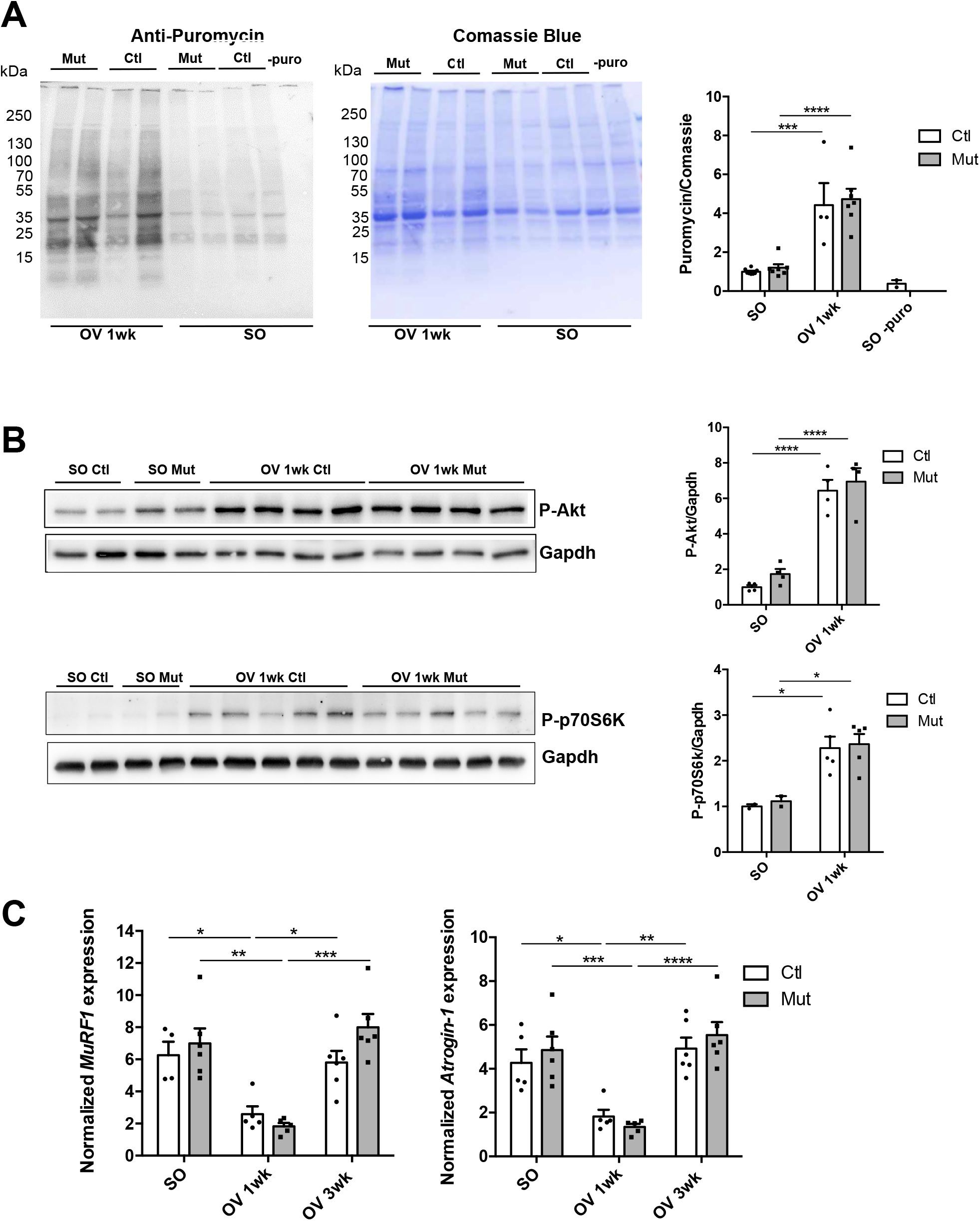
RhoA is not required for Akt dependent signaling and protein synthesis. **(A)** Representative SUnSET experiment performed on Ctl and Mut plantaris muscles before (SO) and after 1wk OV,. Puromycin was injected 30 min before harvesting. Puromycin-labeled peptides were quantified by Western blot using anti-Puromycin antibody. Coomassie was used as a loading control (n=4-8). Ratio of the quantification of Puromycin to Coomassie is shown in the right panel. **(B)** Phosphorylated Akt and p70S6K were analysed by Western blot in Ctl and Mut *plantaris* before (SO) and after 1wk OV. Gapdh was used as a loading control (n=4). Ratio of the quantification of P-Akt or P-p70S6K to Gapdh is shown in the right panel. **(C)** *MuRF1* and *Atrogin-1* mRNA expressions were analysed by RT-qPCR in Ctl and Mut *plantaris* before (SO) and after 1 and 3wk OV (n=4-6). Data were normalized by *Hmbs* expression. Data and mean±SEM. *pvalue<0.05, **pvalue<0.01, ***pvalue<0.001, ****pvalue<0.0001

**Figure S2.**
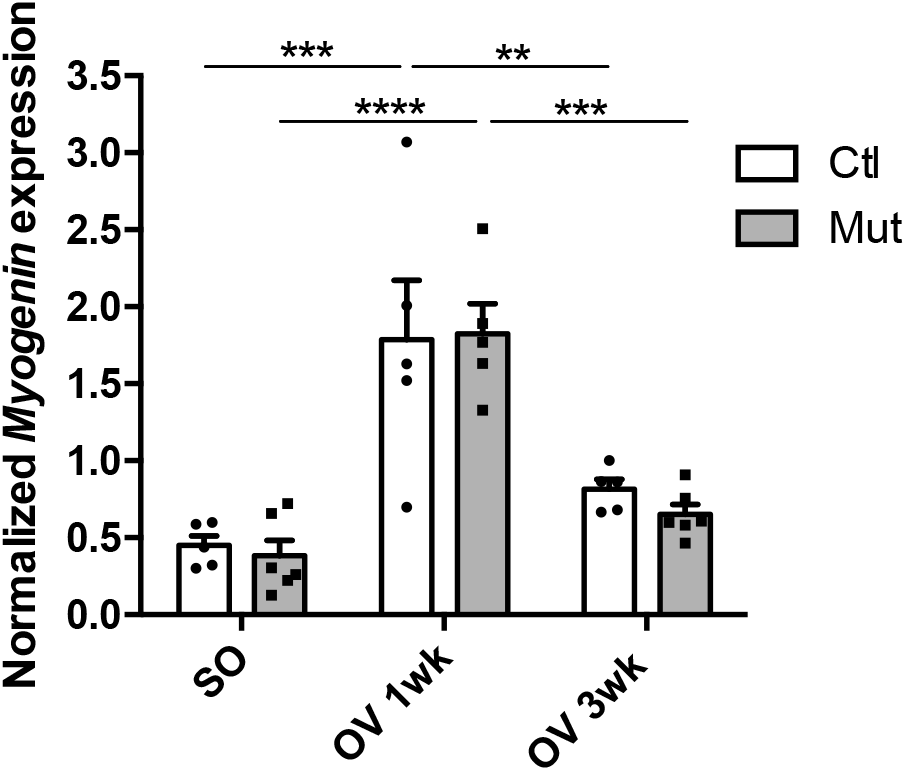
RhoA loss within myofibers does not affect *Myogenin* expression. *Myogenin* mRNA expression was analysed by RT-qPCR in Ctl and Mut *plantaris* muscles before (SO) and after 1 and 3 wk OV (n=5-6). Data were normalized by *Hmbs* expression. Data and mean±SEM. **pvalue<0.01, ***pvalue<0.001, ****pvalue<0.0001.

**Figure S3.**
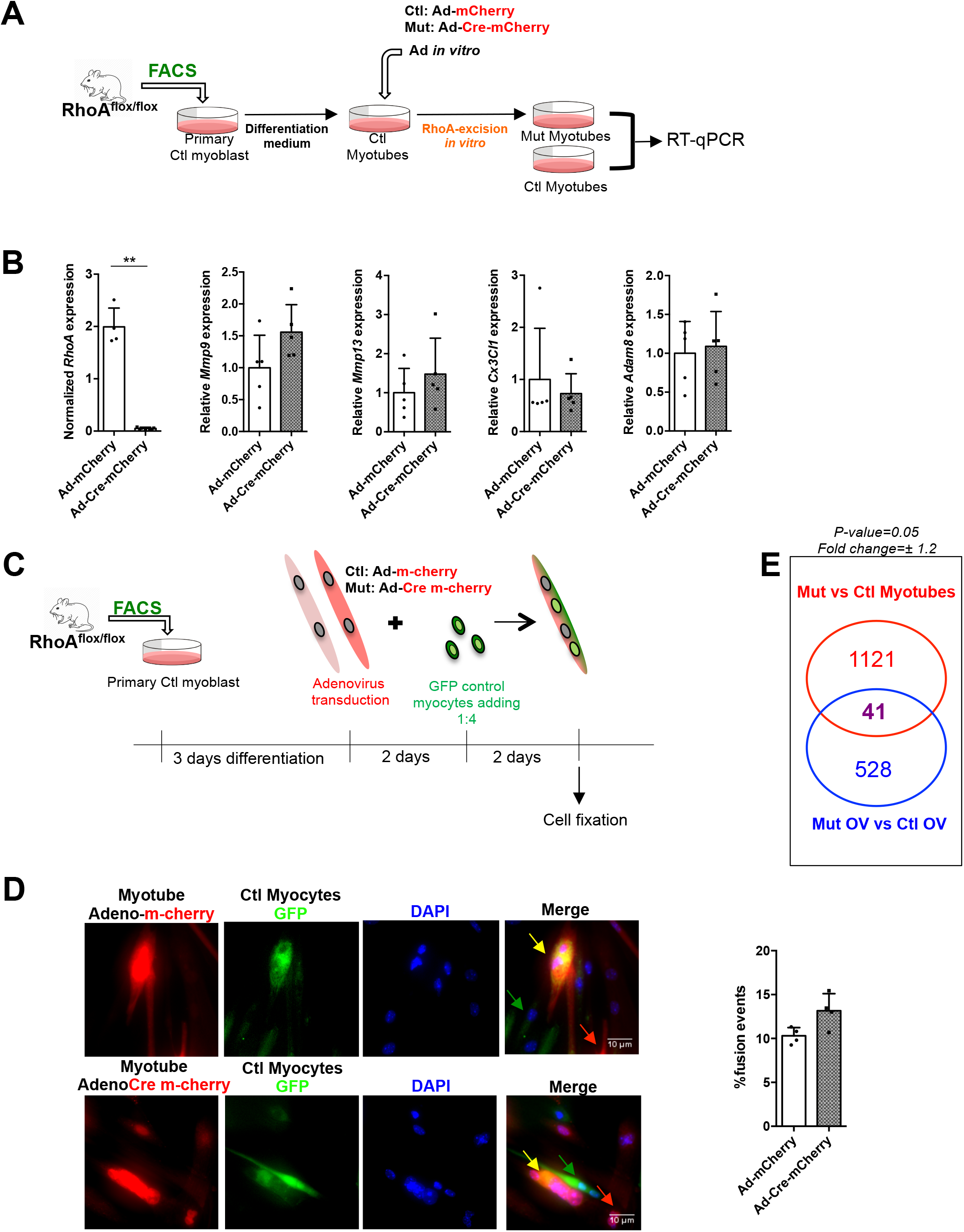
RhoA loss within myotubes *in vitro* does not affect myoblast fusion. **(A)** RhoA^flox/flox^ Myotubes (at 3 days post differentiation) were transduced with Ad-mCherry (Ctl, red) or Ad-Cre-mCherry (Mut, red) and harvested 2 days later. **(B)** *RhoA, Mmp9*, *Mmp13*, *Adam8* and *Cx3Cl1* expressions were analysed by RT-qPCR *in RhoA^flox/flox^* myoblasts transduced with Ad-mCherry or Ad-Cre-mCherry to induce RhoA loss. Data were normalized by *Hmbs* expression (n=4-5). **(C)** Myotubes transduced with Ad-mCherry (Ctl, red) or Ad-Cre-mCherry (Mut, red) were mixed with Ctl myocytes transduced with Lenti-GFP (green). After 48hr of co-culture, myotubes were analysed for dual labeling. **(D)** The percentage of dual-labeled cells per total number of nuclei (fusion events) was scored (n=3). **(E)** Affymetrix analysis has been performed from mRNA from Myotubes transduced with Ad-mCherry (Ctl) or Ad-Cre-mCherry (Mut). Venn diagram showing the intersection between genes differentially regulated by RhoA (fold change>1.2; pvalue<0.05) in Myotubes, and *plantaris* overloaded muscles (Mut OV vs Ctl OV). Data and mean±SEM. ** pvalue<0.01.

**Figure S4.**
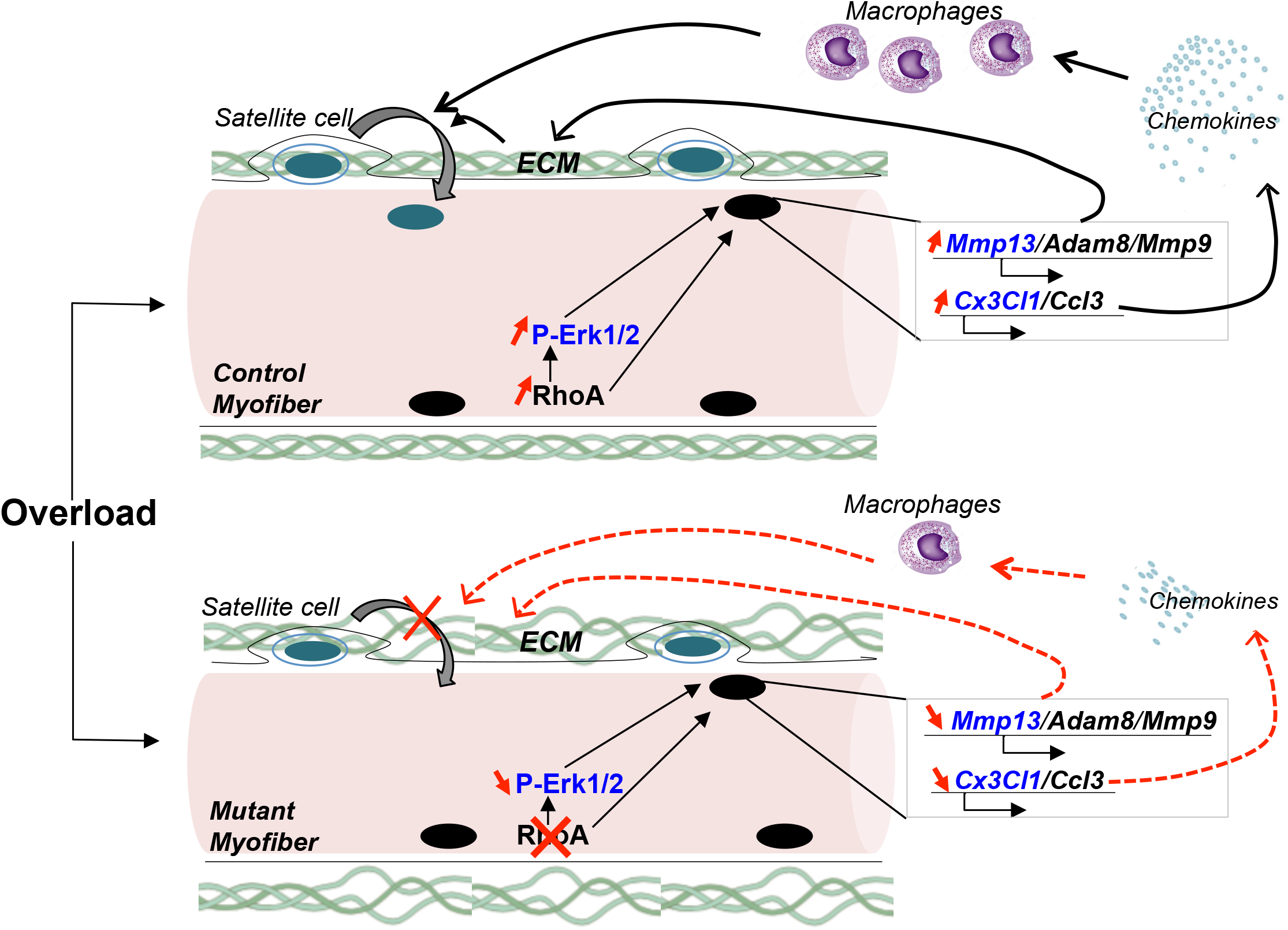
Schematic model: in response to increased workload, RhoA within myofibers gets activated and modulates *Mmp13/Cx3cl1* and *Mmp9/Adam8/Ccl3* expressions in a Erk-dependent and independent manner respectively. In turn, Mmp9/Mmp13/Adam8 modify ECM organization and Cxcl3/Ccl3 chemokines attract macrophages that will remodel SC microenvironment and thus support skeletal muscle hypertrophy and SC accretion.

**Table S1.** Microarray GO analysis in control *plantaris* 1 week after overload versus SO.

| **Canonical pathways** | **-log(pvalue)** | **Z-score** |
| --- | --- | --- |
| Integrin Signaling | 1.07E01 | 4.07 |
| IL-8 Signaling | 8.25E00 | 4.12 |
| Fcγ<sub>3</sub> Receptor-mediated Phagocytosis in Macrophages and Monocytes | 8.18E00 | 5.25 |
| Remodeling of Epithelial Adherens Junctions | 7.69E00 | 2.23 |
| Leukocyte Extravasation Signaling | 7.25E00 | 3.75 |
| Regulation of Actin-based Motility by Rho | 6.92E00 | 4.23 |
| ***RhoA* Signaling** | **6.61E00** | **3.71** |
| Rac Signaling | 6.33E00 | 4.15 |
| Glioma Signaling | 6.09E00 | 2.14 |
| Phospholipase C Signaling | 5.82E00 | 3.16 |
| Actin Nucleation by ARP-WASP Complex | 5.81E00 | 3.88 |
| Cdc42 Signaling | 5.7E00 | 2.54 |
| Dendritic Cell Maturation | 5.37E00 | 3.44 |
| Colorectal Cancer Metastasis Signaling | 5.2E00 | 3.44 |
| Signaling by Rho Family GTPases | 5.17E00 | 3.47 |
| p70S6K Signaling | 5.15E00 | 2.56 |

**Table S2.** Common genes differentially expressed in Myotubes expressing or not RhoA and in Control vs Mutant overloaded-*plantaris* that were similarly regulated (up or down)

| *Gene Symbol* | **Myotubes lacking RhoA vs Clt** |  | **Overloaded-plantaris Mut vs Clt** |  |
| --- | --- | --- | --- | --- |
|  | *Fold Change* | *p-value* | *Fold Change* | *p-value* |
| AmmerR1 | -1,92 | 0,009 | -1,233 | 0,00182 |
| Bcl2A1 | -1,63 | 0,0051 | -1,623 | 0,0148 |
| Csnk1g1 | -1,76 | 0,004 | -1,235 | 0,00366 |
| Cst12 | -1,33 | 0,0027 | -1,221 | 0,00989 |
| Lrp11 | -1,91 | 0,0082 | -1,25 | 0,022 |
| RhoA | -1,68 | 0,0181 | -1,204 | 0,00418 |
| Slx4 | -1,43 | 0,0054 | -1,283 | 0,0117 |
| Znf8 | -1,44 | 0,0144 | -1,216 | 0,00272 |
| 1700001F09Rik (includes others) | 1,28 | 0,0481 | 1,306 | 0,0171 |
| Akap1 | 2,78 | 0,0262 | 1,25 | 0,031 |
| Arpc5l | 1,46 | 0,0229 | 1,358 | 0,0146 |
| Chchd1 | 1,35 | 0,0414 | 1,353 | 0,0218 |
| Cog4 | 2,17 | 0,0284 | 1,267 | 0,0121 |
| Cox19 | 1,69 | 0,0458 | 1,263 | 0,00886 |
| Crygd | 1,48 | 0,0129 | 1,369 | 0,021 |
| Lonp1 | 1,5 | 0,044 | 1,246 | 0,0104 |
| Mrpl12 | 1,58 | 0,0065 | 1,219 | 0,000396 |
| Mrpl53 | 1,72 | 0,0404 | 1,248 | 0,0032 |
| Ndufb5 | 1,23 | 0,0274 | 1,21 | 0,00386 |
| Neurl2 | 1,81 | 0,0152 | 1,271 | 0,00937 |
| Pipox | 1,24 | 0,0384 | 1,267 | 0,0319 |
| PolR2F | 2,32 | 0,0366 | 1,302 | 0,00096 |
| Ppa1 | 2,56 | 0,0391 | 1,226 | 0,0141 |
| Timm10 | 1,86 | 0,0108 | 1,217 | 0,0458 |
| Twf2 | 2,38 | 0,014 | 1,26 | 0,00321 |

**Table S3.**
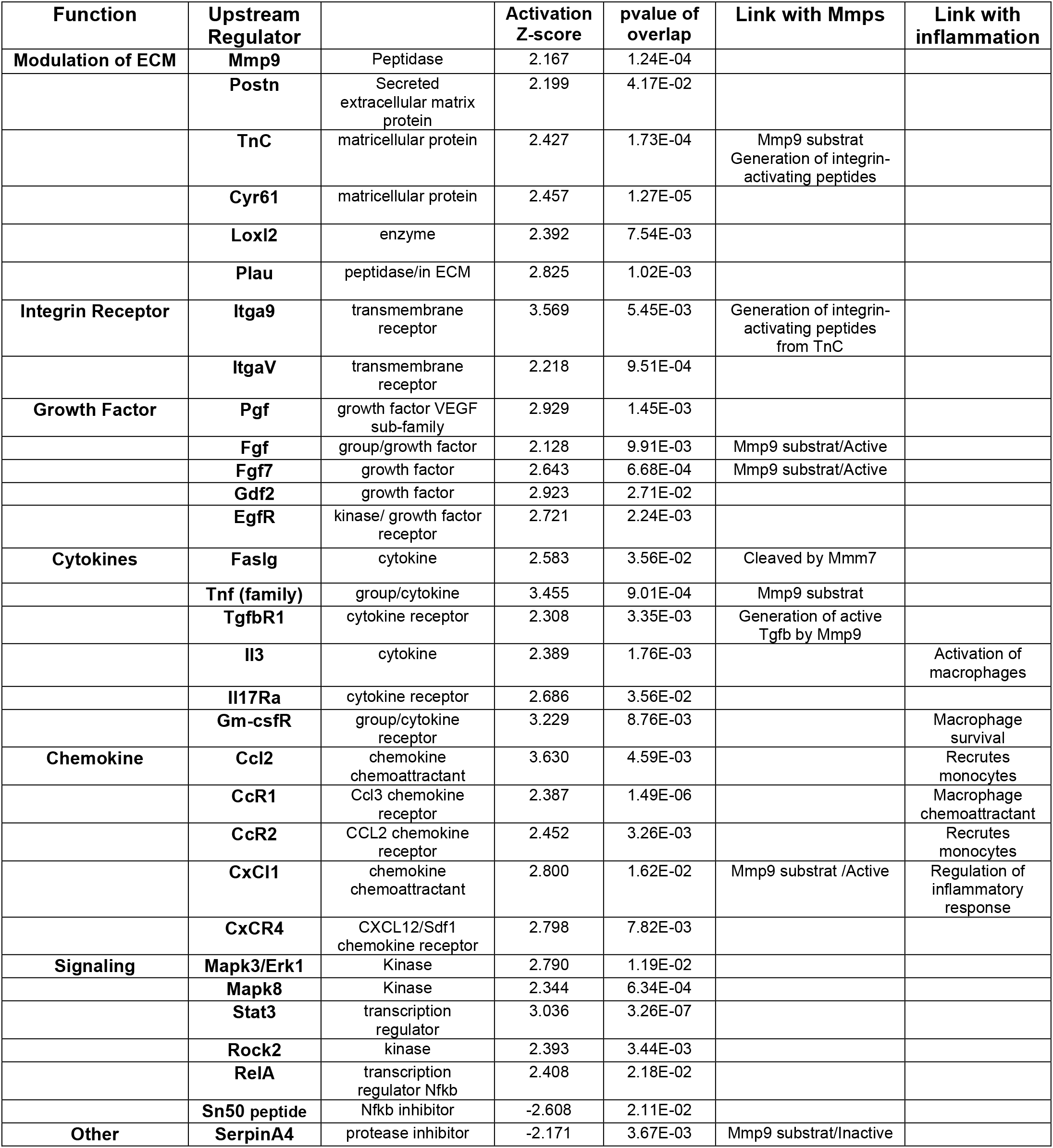
Predicted activated/inhibited Upstream Regulators in Control *plantaris* 1 week after overload vs SO.

